# The sources and transmission routes of microbial populations throughout a meat processing facility

**DOI:** 10.1101/758730

**Authors:** Benjamin Zwirzitz, Stefanie U. Wetzels, Emmanuel D. Dixon, Beatrix Stessl, Andreas Zaiser, Isabel Rabanser, Sarah Thalguter, Beate Pinior, Franz-Ferdinand Roch, Cameron Strachan, Jürgen Zhangellini, Monika Dzieciol, Martin Wagner, Evelyne Mann

## Abstract

Microbial food spoilage is responsible for a considerable amount of waste and can cause food-borne diseases in humans, particularly in immunocompromised individuals and children. Therefore, preventing microbial food spoilage is a major concern for health authorities, regulators, consumers, and the food industry. However, the contamination of food products is difficult to control because there are several potential sources during production, processing, storage, distribution, and consumption, where microorganisms come in contact with the product. Here, we conduct the first study that uses high-throughput full-length 16S rRNA gene sequencing to provide novel insights into bacterial community structure throughout a pork processing plant. Specifically, we investigated what proportion of bacteria on meat are not animal-associated and are therefore transferred during cutting via personnel, equipment, machines, or the slaughter environment. We then created a facility-specific transmission map of bacterial flow which revealed previously unknown sources of bacterial contamination. This allowed us to pinpoint specific taxa to particular environmental sources and provide the facility with essential information for targeted disinfection. For example, *Moraxella* spp., a prominent meat spoilage organism which was one of the most abundant amplicon sequence variants (ASVs) detected on the meat, was most likely transferred from the gloves of employees, a railing at the classification step, and the polishing tunnel whips. Finally, we provide evidence that 1000 sequences per sample provides a reasonable sequencing depth for microbial source tracking in food processing, suggesting that this approach could be implemented in regular monitoring systems.

## Background

As the world population is expected to rise to 9.8 billion by 2050, the global demand for food will increase by approximately 70 % in order to satisfy human needs. Resolving this issue, while also reducing greenhouse gas emissions and protecting valuable ecosystems is one of the greatest challenges of our era. Food security experts estimate that 46 % of the required additional food demand can be achieved by increasing food production, whereas the remaining proportion needs to be attained through sustaining the productive capacity (34 %) and better food demand management (20 %) (Keating et al. 2014). One major element in the reduction of food loss is the prevention of microbial spoilage of food, which is estimated to account for one-fourth of global food waste (Huis In’t Veld 1996). In addition, microbial spoilage of food also poses a key public health concern. The World Health Organization (WHO) reported 600 million foodborne illnesses causing 420.000 deaths worldwide in 2010 alone, indicating that the global burden of food-borne diseases is comparable to those of major infectious diseases (Havelaar et al. 2015). Indeed, the food industry faces major and continuing challenges in trying to lower the extent to which food products become contaminated with pathogenic or spoilage bacteria during primary processing. This is especially true for animal-derived products like poultry, eggs, milk, and pork, which are main vehicles of food-borne diseases (EFSA and ECDC 2018). Thus, microbial meat spoilage is a global health and economic challenge, yet little is known about the microbial diversity in slaughterhouses and meat cutting plants. Pork provides an ecosystem with high water content and nutrient availability for diverse microorganisms, particularly psychrotolerant organisms, which can then grow during storage and lead to spoilage and a reduced shelf life (Gill 1983). The formation of biofilms on processing equipment is also of great concern and can lead to continuous dispersal of microorganisms on the meat (Giaouris et al. 2014). To date, transmission routes of microorganisms during meat processing have been difficult to track and monitor and therefor remained largely elusive (Choi et al. 2013; Sheridan 1998). The main reason for this is that the meat industry is still relying on ISO reference methods applying microbiological techniques to monitor hygiene aerobic colony counts (ISO 4833) and *Enterobacteriacae* (ISO 21528-2) (International Organization for Standardization 2015). Indeed, culture-dependent techniques can be useful in determining the overall hygiene status of a facility, but they fail in describing complex microbial communities and population flows (Nocker et al. 2007). Just recently, first efforts have been made to characterize the microbiome of food processing environments with next-generation sequencing techniques, demonstrating the suitability of these tools to map microbial ecosystems in different sectors of food industry (Bokulich et al. 2015; Hultman et al. 2015; Chaillou et al. 2015; Yang et al. 2016). Based on a previous study, we hypothesized, that about one-third of bacteria on meat are not animal-associated and therefor have to be transmitted during cutting via personnel, from the equipment, or from the machine and slaughter environment (Mann et al. 2016). Using full-length 16S rRNA gene sequencing we were able to identify key carrier points of bacterial contamination and create a facility-specific transmission map of bacterial flows that exposed unique transmission patterns for individual taxa. Although this study primarily aids to optimize slaughter processes to systematically avoid contamination of microbes throughout meat processing, the techniques we applied can also be extended to other food-processing environments. In this way, we can expand our knowledge about microbial transmission routes, thereby improving hygiene standards in food-related industry to increase food safety while minimizing food waste.

## Materials and Methods

### Facility structure and sampling

Samples for this study were taken from an Austrian slaughterhouse with a capacity of 200-250 pigs per hour. Figure 1 illustrates the structural design of the production chain and the sampling positions. *Ante mortem* inspection took place during the offload and holding. The pigs were herded through a series of gates in small groups until they were finally presented for stunning one by one. After electrocution to the head, each pig’s main throat vasculature was cut with a knife and the bodies hanged up to exsanguinate (Position Sticking). Then the carcasses were scalded for three minutes by steam condensation, after which the claw shoes were manually removed. Before the carcasses entered the singeing tunnel for approximately six seconds, they were dehaired by rotating whips and manually pre-singed. Polishing was then performed in another tunnel with rotating whips and water spray (Position Polishing). Eyes and external ear canal were removed from the bodies as they were moved from the “dirty area” to the “clean area”. Operators were not allowed to move from one area to the other. To avoid contaminations from feces, the rectum was sealed off with a small plastic bag. Evisceration was the first step in the clean area, followed by splitting the carcasses in halves with a saw, post mortem inspection, removal of the spinal cord with an aspirator, and classification. After classification, the carcasses passed a shock shower with 4 °C cold water and entered the cooling chamber where they were held for up to 16 hours at 7 °C. The chilled carcasses were then transported on a refrigerated truck to a meat cutting facility.

**Figure 1:**
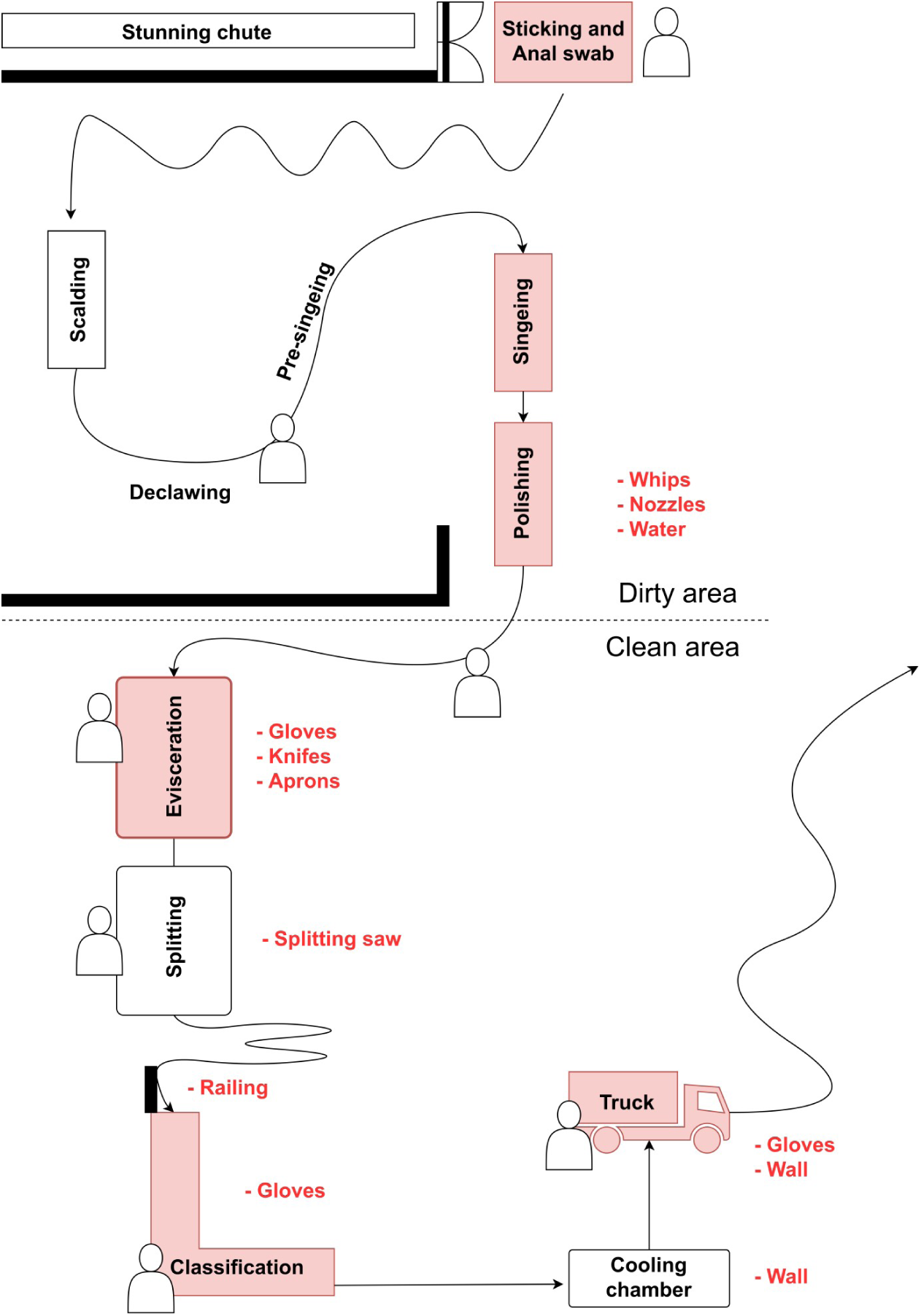
Schematic map of the slaughterhouse depicting the whole processing line. Red shaded boxes indicate locations where skin/meat was sampled. Red text specifies environmental samples taken at corresponding positions. Human sketches point out non-automated locations with working employees.

Twelve pigs from three different farms (four pigs each) were used for this study. Carcass and environmental samples were taken with sterile polyurethane sponges (SampleRight™ Sponge Sampler, World Bioproducts, Woodinville, USA), which recover significantly higher amounts of bacteria compared to sponges made from cellulose, reaching a similar recovery rate as excision methods (Pearce and Bolton 2005). The sampling area was 100 cm^2^ at the back of the carcasses. The back was chosen based on the results of a preliminary test already conducted prior the start of this study, which showed that samples taken from the back harbour similar levels of relevant microorganisms compared to samples taken from the belly or the musculature along the cutting area (Zwirzitz et al. 2019). In total, 84 swabs were taken from carcass surfaces at different processing positions. The carcasses chosen for sampling were ear tagged and followed throughout the entire processing chain so that the same carcasses could be sampled at each position. In addition, 75 swabs were taken from equipment, staff, and infrastructure of the facility. The sampling was done over a period of one day. For a detailed list of samples and sampling positions see supplementary file 1. For sampling, the sponge was swabbed for ten seconds horizontally, then flipped and swabbed again for ten seconds in vertical direction. Then, the sponge was placed back into the sterile plastic bag, which was sealed and chilled in a container placed in the cooling chamber of the facility (4 °C) until sampling was finished. A fresh polyurethane sponge, a new sterile template of 100 cm^2^ and new gloves were used for each new sample. All the samples were transported back to the lab on ice (Transport time: 2 hours). Back in the laboratory, each sponge was squeezed thoroughly, and the obtained liquid was split into two 15 ml falcon tubes, one of which was directly used for cultivation experiments, while the other one was stored at −20 °C until further processing (Molecular analysis).

### Microbiological investigation

The enumeration of aerobic mesophilic counts (AMC) (ISO 4833-2:2013), *Enterobacteriaceae* (EB) (ISO 21528-2:2017) and *Pseudomonadaceae* (PS) was performed after preparing a ten-fold serial dilution in buffered peptone water (BPW) (Thermo Fisher Scientific Inc., Oxoid Ltd., Basingstoke, United Kingdom) up to dilution −10^8^. The dilutions were plated in duplicates on Plate Count Agar (PCA, Thermo Fisher Scientific Inc., Oxoid Ltd.), Violet Red Bile Glucose (VRBG, Thermo Fisher Scientific Inc., Oxoid Ltd.) and Glutamate Starch Phenol Red Agar (GSP, Merck KGaA; Darmstadt, Germany) by surface plating technique and incubated for a maximum of 72 and 48 h at 30, 37 and 25°C, respectively. To determine the AMC/EB and PS counts/cm^2^, microbial colonies between 10 and 300 colony forming units (CFU) were included in the calculation. Presumptive EB and PS isolates were confirmed by Oxidase reaction and biochemical profiling (API 20E, bioMérieux Marcy-l’Étoile, France).

### DNA-extraction, qPCR, and 16S rRNA gene sequencing

In order to increase microbial cell density, samples were centrifuged at 3.220 × rcf for 20 minutes and the pellet was resuspended in 400 µl of 1 × phosphate buffered saline (PBS). The DNA was then extracted from 200 µl with the QIAamp DNA Stool Mini Kit (Qiagen GmbH, Hilden, Germany) according to manufacturer instructions. The elution step of the protocol was modified; instead of 200 µl AE buffer, two times 25 µl DEPC treated water was used. Negative controls (DEPC treated water), one for each used kit, were also extracted together with the regular samples. The DNA concentration of the samples was measured with the Qubit dsDNA HS Assay Kit and Qubit 2.0 Fluorometer (Invitrogen, Thermo Fisher Scientific, Oregon, USA).

The 16S rRNA gene was amplified by qPCR to enumerate total bacterial cell equivalents (BCE) as previously described (Zwirzitz et al. 2019; Supplementary file 2). All qPCR samples and standards were run in duplicates and negative controls were included in each run (Mx3000P qPCR thermocycler, analyzed with MxPro v.4.10 (Stratagene, San Diego, USA)). Total BCE were extrapolated with an average of four 16S rRNA gene copies as estimated by rrnDB, a database for ribosomal RNA operon variation in bacteria and archaea (Stoddard et al. 2015; Větrovský and Baldrian 2013). Statistics for qPCR and plate count data were tested for normal distribution using the Shapiro-Wilk normality test and with visual assessment of qqplots and histograms. The not normal distributed groups were tested using the Wilcoxon-test for connected samples. We applied Benjamini-Hochberg procedure to reduce the false discovery rate. Data are considered significant at p≤0.05.

Amplicon library generation, quality control, and sequencing were performed at the Vienna Biocenter Core Facilities NGS Unit (www.vbcf.ac.at). Full-length 16S rRNA gene libraries of 133 samples (including 3 negative controls) were prepared using bacteria specific primers 27F (5’-AGRGTTYGATYMTGGCTCAG-3’) and 1492R (5’-RGYTACCTTGTTACGACTT-3’). Barcodes were added during a second round of amplification with Pacbio Barcoded Universal primers, so that the amplicons could be multiplexed on three SMRT cells. Sequencing was carried out on a Pacbio Sequel machine with 2.1 chemistry. Detailed library preparation and sequencing procedure is available online https://www.pacb.com/wp-content/uploads/Procedure-Checklist-Full-Length-16S-Amplification-SMRTbell-Library-Preparation-and-Sequencing.pdf. Each SMRT cell generated approximately 50 GB of raw data producing 641.939 sequences/cell on average.

In addition to the full-length 16S rRNA gene of 133 samples sequenced on a Pacbio Sequel machine, the V3-V4 region of the 16S rRNA gene from 52 of these samples was also sequenced on an Illumina MiSeq sequencing platform with a 300 bp paired-end read protocol. The PCR reactions were performed as described in Klindworth *et al*. using the forward primer 341f (5’-TCGTCGGCAGCGTCAGATGTGTATAAGAGACAG) and the reverse primer 785r (5’-GTCTCGTGGGCTCGGAGATGTGTATAAGAGACAG) (Klindworth et al. 2013).

### Sequence processing and analysis

Accurate full-length 16S rRNA gene sequences were generated using Pacbio’s single-molecule circular consensus sequencing. The circular consensus reads (ccs) were determined with a minimum predicted accuracy of 0.99 and the minimum number of passes set to 3 in the SMRT Link software package 5.1 (“Pacific Biosciences SMRT® Tools Reference Guide.” 2018). After demultiplexing, the ccs were further processed with DADA2 (version 1.9.1) to obtain amplicons with single-nucleotide resolution (Callahan et al. 2016, 2018). Similarly, sequences generated on Illumina’s MiSeq platform were also processed with DADA2 and equivalent parameters in order to achieve a maximum comparability between the two datasets. Detailed parameters of our sequence processing workflow in the form of a reproducible R markdown document can be found in the supplements (Supplementary file 3). Amplicon sequence variants (ASVs) were assigned a taxonomy using a DADA2 formatted version of the genome taxonomy database release 03-RS86 (Alishum 2019; Parks et al. 2018). After initial quality filtering, samples with less than 200 reads and ASVs with less than 5 reads were removed. Additionally, contaminant ASVs were detected and removed with the R package “*decontam*” using a prevalence-based contaminant identification with a p-value cutoff of 0.5 (Davis et al. 2018). Microbial community analysis was performed within the “*phyloseq*” and “*tsnemicrobiota*” packages and visualized with ggplot2 in R (McMurdie and Holmes 2013; Wickham 2016; Lindstrom 2017). The ecological origin of ASVs was inferred by an extensive literature search for ASVs with a cultured representative (1-2 articles that describe its isolation) and by checking the isolation source in GenBank for sequences assigned to uncultured species described from environmental samples. Each ASV could have more than one possible origin if multiple different sources were specified. The supposed ecological provenance was then estimated by counting occurrences of the top 500 ASVs in certain habitats.

Microbial source tracking was achieved with the software SourceTracker (version 1.0.0) and default parameters (Knights et al. 2011). Samples taken from the facility environment and from the skin of the animals were assigned as sources whereas meat samples were assigned as sinks. The Illumina dataset (min: 7.712; mean: 19.119; max: 30.340 sequences per sample) was rarefied to 7.712, 5.000, 1.000, 500, and 200 reads per sample prior to SourceTracker analysis in order to infer the influence of sequencing depth on the performance of SourceTracker. To investigate the variation of same sized datasets this procedure was repeated three times for the larger datasets and ten times for the dataset with 200 sequences, because we expected a higher variance when using 200 random sequences. For each dataset size the goodness of fit was determined. In detail, the squared difference between randomly generated datasets and the original dataset (i.e. including all sequences) was calculated and the match rate for both, rarefied and original dataset were checked for correlation with spearman correlation coefficient and correlation plots. The normal distribution of the squared differences for each dataset size was investigated by using the Shapiro-Wilks test and the homoscedasticity was analysed by applying the Levene test. Due to non-normal distribution of the data, a Kruskal-Wallis rank sum test, followed by a Dunn test with Benjamini-Hochberg alpha-adjustment for post-hoc analysis was applied in order to compare the goodness of fit of the different dataset sizes. Additionally, the hit ratio of each dataset size was calculated. The hit ratio is defined as percentage of correctly assigned contamination sources (including ASVs with an unknown source), whereby the original dataset was used as reference. Further a comparison of amounts of unknown classified ASVs (unknown classification rate) was accomplished. Due to significant differences between the production steps regarding to the correctly assigned samples in the unknown sources (data not shown), the differences between the amounts of unknown classified ASVs for each randomly generated dataset and the original dataset were separately calculated. The significant differences between the hit ratios and the rate of unknown classified ASVs were also analysed with Kruskal-Wallis rank sum test and Dunn test for post-hoc analysis. All statistical analysis was performed in R.

## Results

### Bacterial cell counts strongly differ between sampling positions and surface locations

In order to get an initial basic understanding of the microbiological status along the processing line of the facility we determined total bacterial cell equivalents using 16S rRNA gene qPCR, as well as AMC, *EB*, and *PS* counts by applying ISO reference methods (ISO 4833, ISO 21528-2) (Figure 2). The animals entered the facility with a high microbial load on skin (Figure 2, “Sticking”), which was reduced significantly after singeing. In the next step (“Polishing”), AMC and *PS* counts increased significantly, while *EB* counts were not detected until that point, but were found at low levels at the evisceration and classification step and in environmental samples (Gloves, knifes, aprons, etc.). In general, aerobic counts mesophilic counts along the slaughter line ranged from 4.14 × 10^3^ CFU/cm^2^ after singeing to 5.21 × 10^6^ CFU/cm^2^ at sicking. The highest levels of bacteria in the environment were found at the whips of the polishing tunnel (2.19 × 10^7^ CFU/cm^2^). BCE counts ranged from 1.64 × 10^2^ BCE/cm^2^ after singeing to 1.20 × 10^5^ BCE/cm^2^ at sticking and were not significantly higher after polishing.

**Figure 2:**
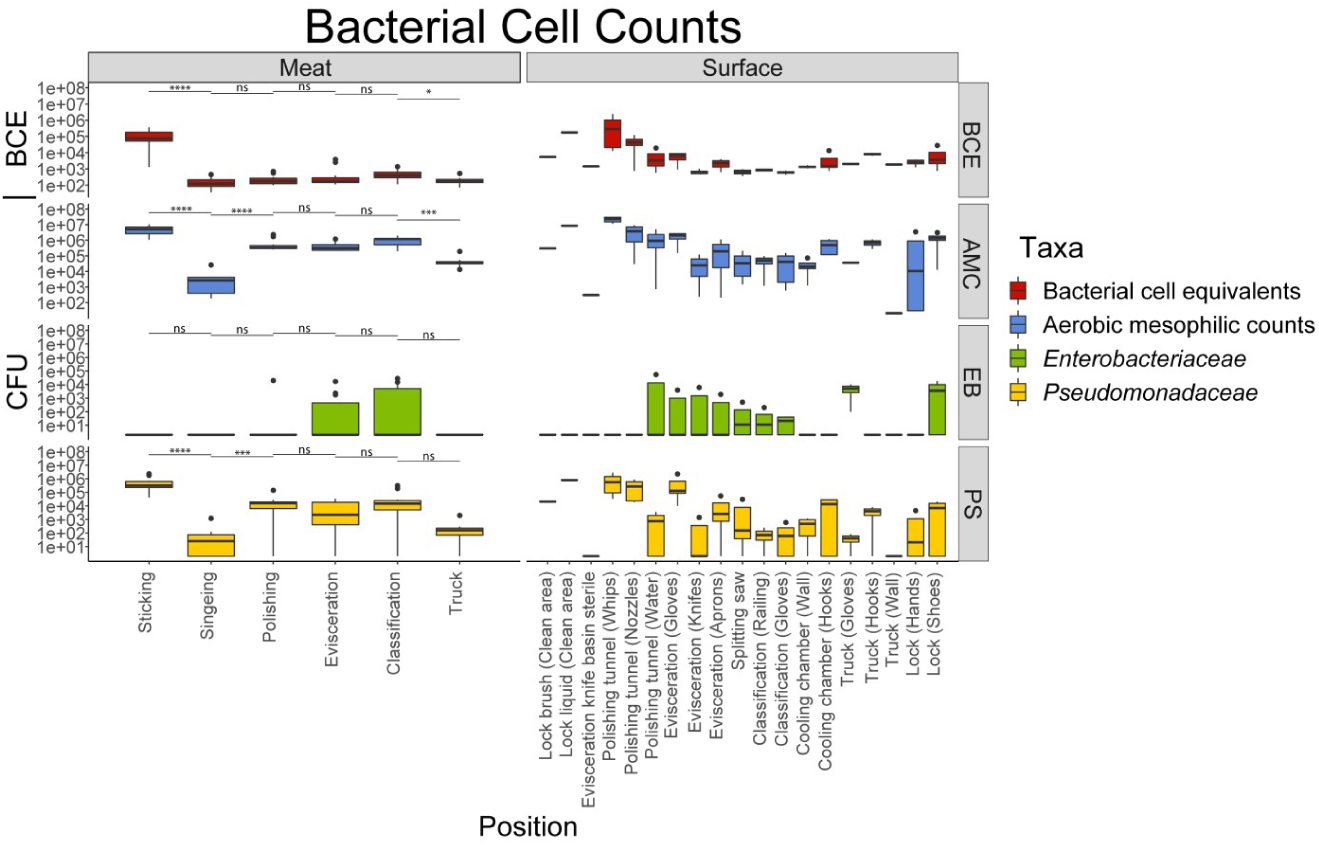
Bacterial cell equivalents (BCE) and colony forming units (CFU) of aerobic mesophilic counts (AMC), Enterobacteriaceae (EB), and Pseudomonadaceae (PS) for the different sampling positions. Levels of significance: ns: p>0.05, *: p≤0.05, **: p≤0.01, ***: p≤0.001.

### Microbial community structure changes throughout the processing line

On average, 1186 high quality full-length 16S rRNA gene sequences per sample remained after stringent quality filtering. A rarefaction curve indicated that most, but not all samples were sequenced deep enough to infer the full diversity of microorganisms in the samples (Supplementary file 4). All analysed alpha diversity indices showed a similar pattern, where there is an overall decrease of microbial species diversity from start to end of the processing line, but with a transient increase after the polishing step (Figure 3A). Beta diversity analysis revealed two major shifts in the microbial community structure (Figure 3B). The differences in microbial composition between samples from the skin and rectum (Positions “Sticking” and “Anal Swab”) of the pigs was higher compared to samples later in the processing line. A shift occurred during the singeing step which is also reflected by significantly reduced microbial numbers as well as species diversity. From that point on the microbial community stayed relatively constant until the end of the processing line between the stations “Classification” and “Truck” at which the communities dispersed from each other.

**Figure 3:**
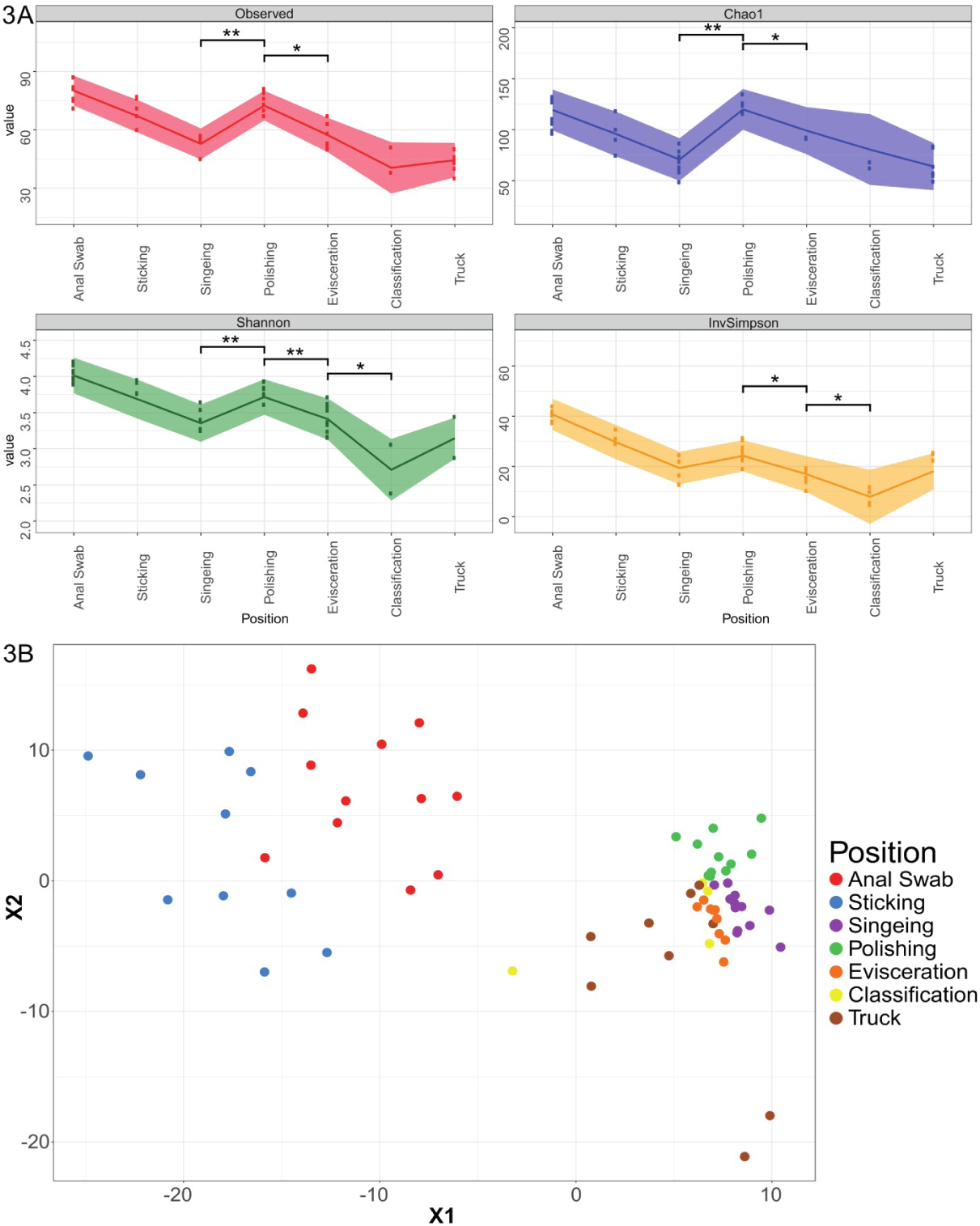
**(A):** Change in alpha diversity indices of meat samples over time. Points represent individual samples, the trend lines connect the means, and shaded regions indicate the standard error. Levels of significance: *: p≤0.05, **: p≤0.01. **(B):** t-SNE plot of Bray-Curtis distances based on 16S rRNA gene libraries. Each point represents values from individual libraries with colors expressing meat samples from different positions along the processing line.

A majority (91%) of the full-length 16S rRNA gene sequences could be assigned to specific genera and 74% to the species level. The 50 most abundant ASVs show heterogeneous distributions and relative abundances across all meat samples (Figure 4). The shift in the microbial community structure that was observed in beta diversity analysis can also be discerned by the relative abundances of ASVs. At the beginning of the slaughterline (Anal Swab and Sticking station), the 50 most abundant ASVs make up less than 20% of the total community and have higher levels of *Helicobacter*, and *Curvibacter*, compared to samples from the other positions. In contrast, the genera *Anoxybacillus, Chryseobacterium*, and *Moraxella* were the most abundant ASVs in the meat samples after the singeing step, making up more than 50% of the sequences. The same ASVs that were highly abundant on the meat samples were also frequently detected in the surface samples of the facility environment. However, the distribution of these ASVs throughout the facility was varied in terms of their location specificity and prevalence. For instance, ASVs belonging to the genus *Bacillus*_S or *Moraxella* were homogenously distributed across many different positions, whereas others were detected only at very specific locations e.g. *Luteimonas_A* and *Helicobacter_F* at the polishing tunnel or W16RD (NCBI taxonomy: *Sphingomonas*) at the wall of the cooling chamber (Figure 4 and Supplementary file 5).

**Figure 4:**
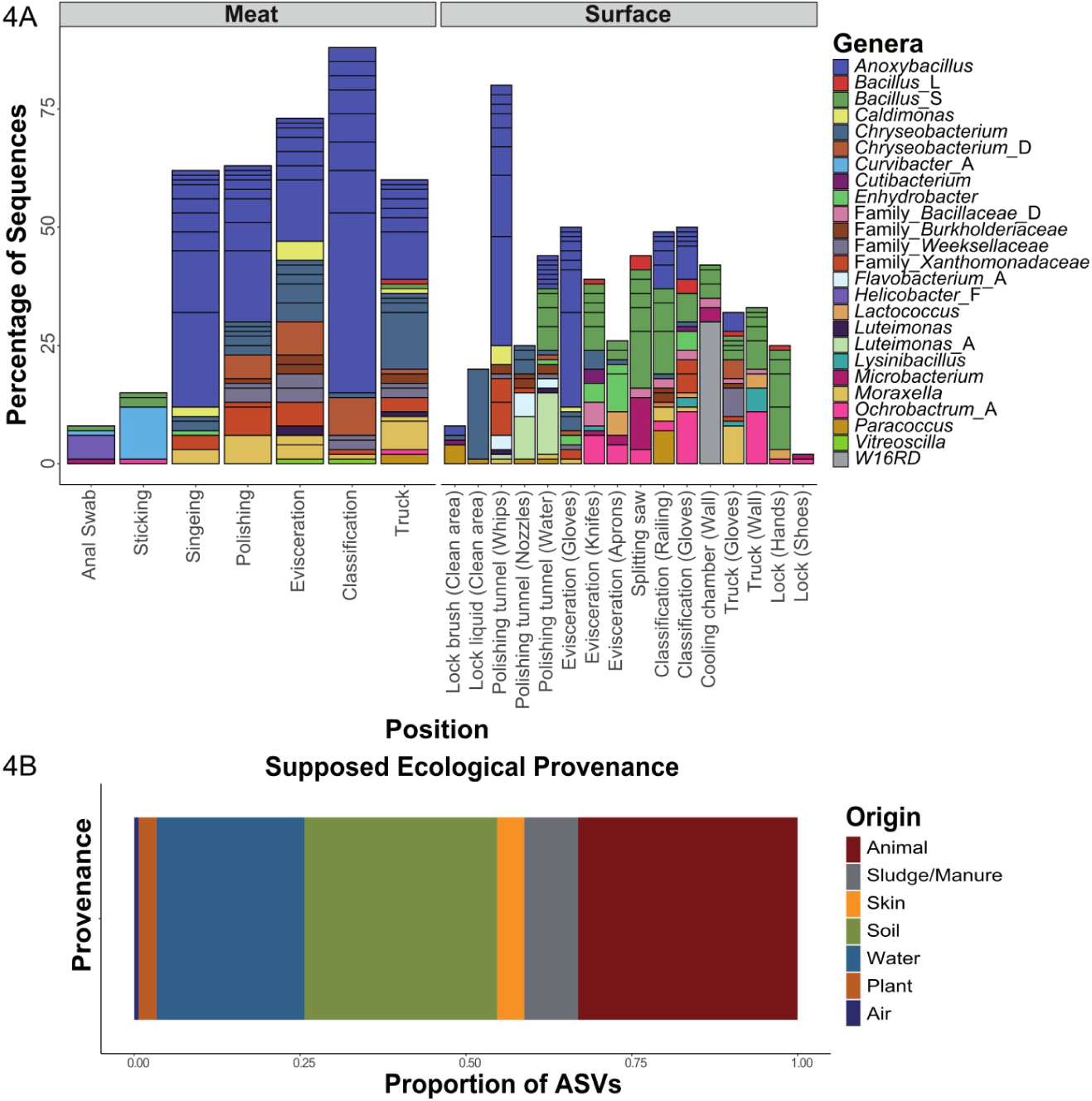
**(A):** Genus-level classification of the 50 most abundant ASVs parted by type (Meat or Surface). Data represents average of ASV counts from replicate libraries for each category. Individual ASVs are separated by a black line within the bar graph. ASVs assigned to candidate genera that do not have a name assignment yet are indicated with “Family_”. Genera names with an alphabetic suffix indicate genera that are polyphyletic and were therefor subdivided in the genome taxonomy database. **(B):** Distribution of the supposed ecological provenance of the 500 most abundant ASVs.

### Variances in spatial distribution of microorganisms throughout the facility result in distinct transmission routes for specific taxa

Transfer of microorganisms from environmental samples to meat samples was inferred using the software SourceTracker. Samples collected from pork carcasses were designated as sinks for testing against the communities of samples from the anal region, the skin, and from the equipment and material surface (source). Alpha diversity indices had already indicated transfer of new species onto the meat at the polishing step (Figure 3A). This was confirmed by SourceTracker analysis showing a high contribution of polishing tunnel samples (nozzles, water, and especially whips) to the microbial community of meat samples (Figure 5). Anal swab and sticking positions were not identified as one of the major sources of bacterial contamination despite having shown high alpha diversity and bacterial cell count values. Hence, the decontamination processes, e.g. singeing, likely eliminated most of the bacteria that were initially on the pork carcass when pigs entered the facility. Furthermore, the small contribution of anal swab samples validates the effectiveness of the general practice in the analyzed facility to seal off the rectum with a small plastic bag before evisceration to avoid contamination with faecal matter. The evisceration step is usually considered to be a critical point of re-contamination if cutting processes are not executed with high hygiene standards. In this case, transfer of microorganisms from aprons and knifes was low but a high proportion (11.4%) of microorganisms detected on meat samples taken from the last position (Truck) originated from the gloves of employees performing the evisceration. Gloves of employees were also identified as major contamination sources at other positions (Truck (8.3%), classification (2.8%)). Furthermore, a railing that all carcasses touched while passing the classification contributed to the microbial composition of the meat samples (4.4%). Environmental samples without direct contact to the carcasses (Lock and wall samples) were also not identified as major contamination sources, suggesting that most bacteria are transferred through direct contact with the surface and that air transfer is marginal. One third (31.6%) of the bacteria on meat could not be linked to a specific source and was therefore attributed to an unknown source. Possible reasons for this are that the sequencing depth for some samples was too low or that the primary source of these bacteria was missed during sampling.

**Figure 5:**
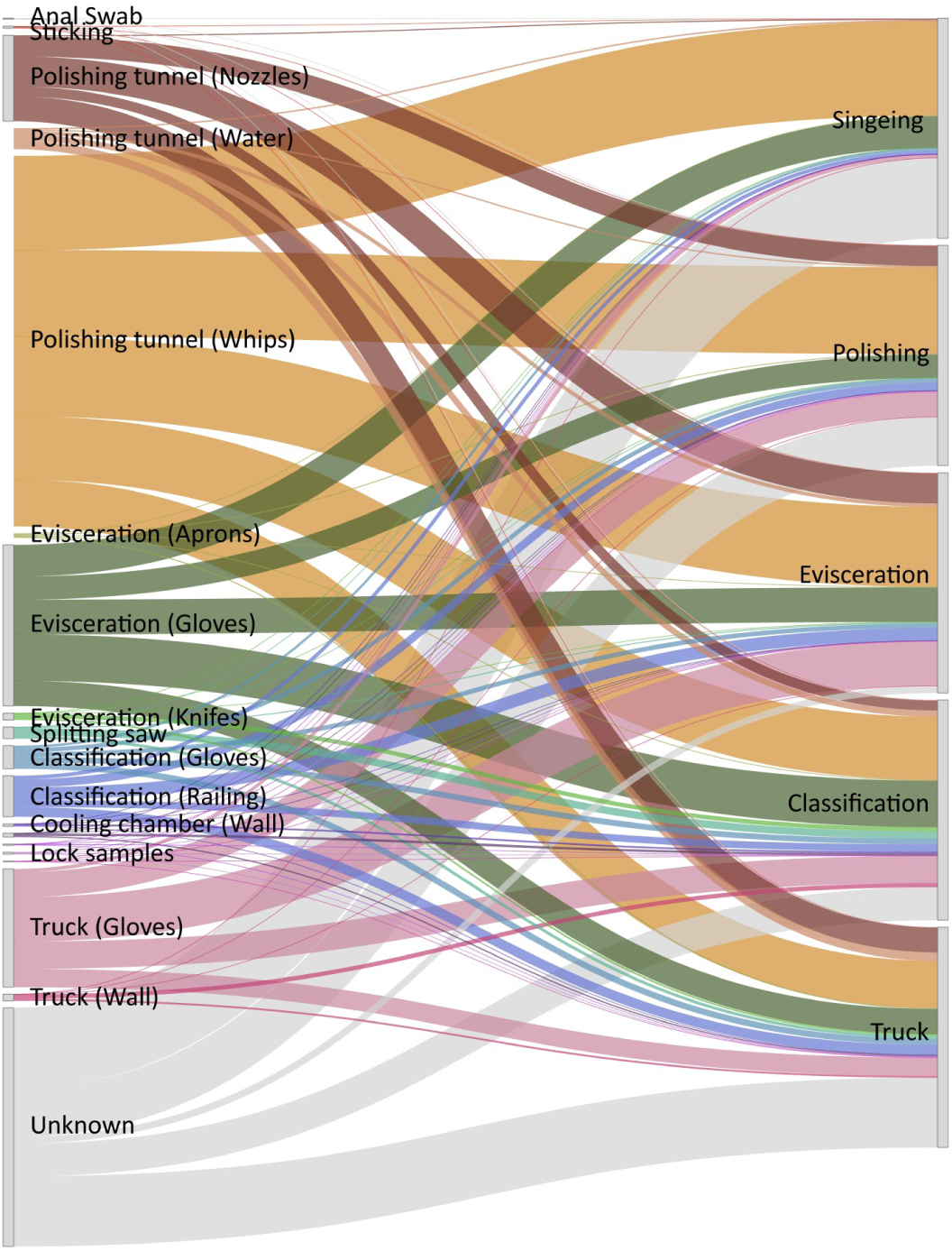
Source environment proportions for meat samples estimated using SourceTracker and visualized as a Sankey flow diagram. Environmental source samples are represented on the left and meat samples, as sinks, are shown on the right. The line width of individual flows between them illustrates the average contribution of microorganisms from source samples to the microbial community of respective sink samples.

Next, we inferred the ecological provenance of the detected microorganisms by an extensive literature search and by checking the isolation source in GenBank. Our initial hypothesis that about one third of the bacteria on meat are not animal-associated but rather originate from the environment was surpassed with this method of evaluation. We estimate that more than half of the 500 most abundant ASVs detected on meat were not originally part of the animals’ microbiota, but were eventually transferred during processing (Figure 4B).

Finally, we were interested if certain microbial species were transmitted from specific sources or if they were spread throughout the whole facility. We investigated the distribution of all detected genera by determining their relative abundances across all meat and environmental positions. This was done for each individual genus (Supplementary file 6 and 7). Many genera were unique to specific sites demonstrating that they occupy particular environmental niches in the facility. In addition, we predicted the relative contribution of genera that are associated with meat spoilage or that include relevant pathogens based on the SourceTracker results (Figure 6). Indeed, some of the microorganisms could be pinpointed to a specific location from where they disseminated, whereas others were found to have multiple possible origins. For example, the genus *Escherichia* originated almost exclusively from the anal swab samples. On the other hand, *Lactococcus, Staphylococcus, Chryseobacterium*, and *Moraxella* species were predicted to be transmitted from various different positions. In general, there was little clustering of taxa based on shared-source similarity in the heatmap, supporting the observation that most analysed taxa have a unique transmission pattern. Some microorganisms were predicted to have similar contributing positions, e.g. *Flavobacterium* and *Lactobacillus_*H from the gloves at the evisceration step or *Lactococcus* and *Bacillus*_L which shared the splitting saw as a common source. A closer look at the phylogenetic relationship of ASVs assigned to the polyphyletic genus *Chryseobacterium* revealed that different populations were transferred from respective sources (Supplementary file 8). A strain classified as *Chryseobacterium* sp. Leaf405 in NCBI taxonomy (GCF_001425355.1 in GTDB), was spread across the polishing tunnel and evisceration positions, whereas others have been found exclusively at the polishing tunnel (*Chryseobacterium indoltheticum*) or at the locks (genus *Chryseobacterium*; no species verification). Other prominent meat spoilage microorganisms, for example *Moraxella* spp., which was also one of the most abundant ASVs on the meat, was most likely transferred from the polishing tunnel whips, gloves of employees as well as from the railing at the classification step. Overall, we were able to identify transmission routes for the majority of the genera that include relevant meat spoilage organisms or pathogens. However, the origin of some taxa (e.g. *Fusobacterium_*C and *Pseudomonas*_E) remains elusive since their contribution to the meat microbiota was attributed to an unknown source.

**Figure 6:**
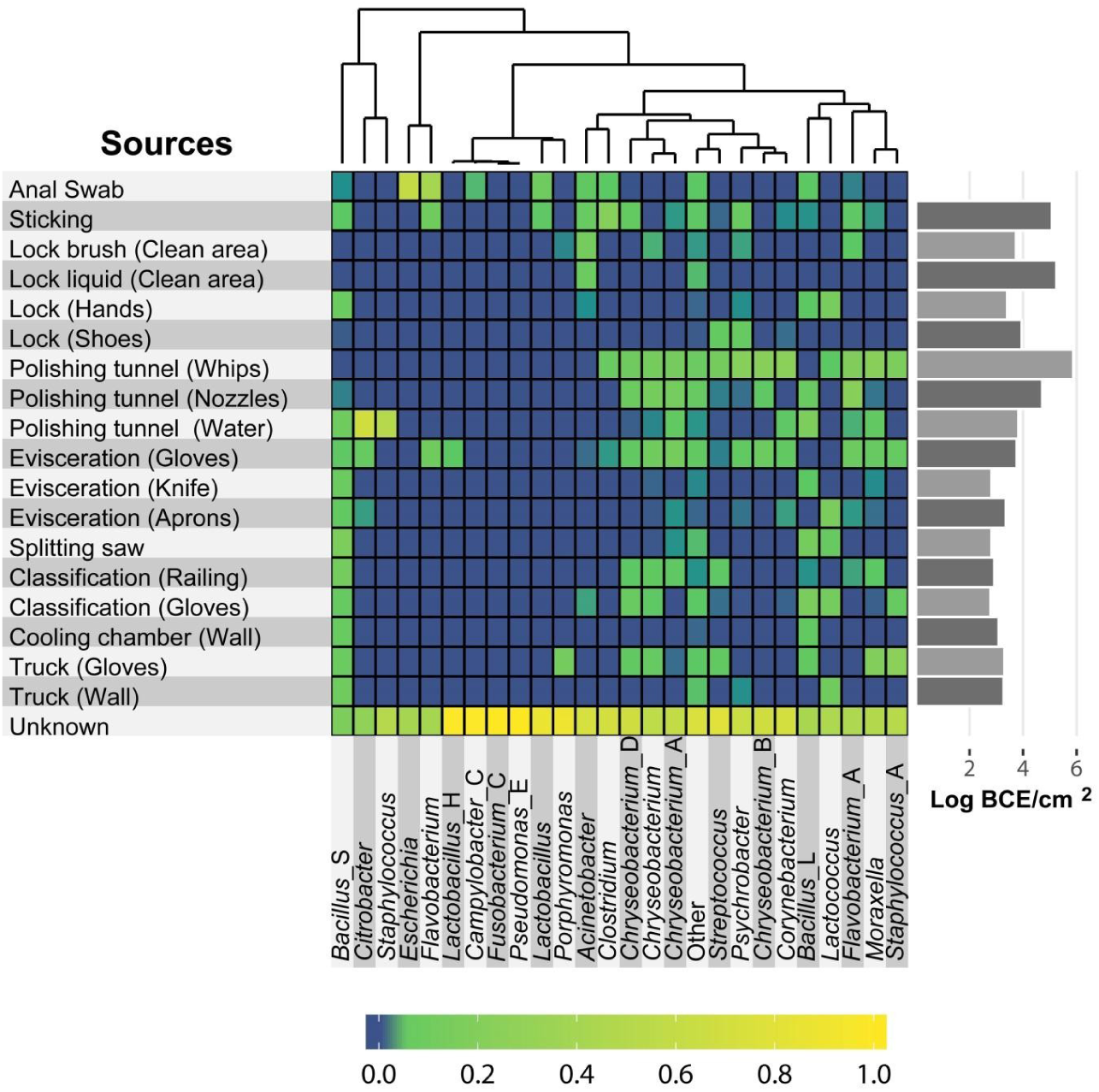
Heatmap showing the predicted relative contribution of specific genera from different source environments. Only taxa that are associated with meat spoilage or include relevant pathogens are shown. Non-relevant taxa were grouped into a single group called “Other”. The relative contribution of each genus (rows) from each source (columns) is indicated by the colour of the intersecting tile. Taxa are clustered by shared-source similarity. BCE=Bacterial cell equivalents as determined by 16S rRNA gene qPCR.

### Sequencing depth has little effect on the performance of SourceTracker

To test whether the shallow sequencing depth (avg.: 1.186 sequences/sample) of the “Pacbio” dataset was sufficient for a comprehensive analysis with SourceTracker, we additionally sequenced a subset of the samples on an Illumina Miseq machine. This dataset was then rarefied to different sizes and SourceTracker analysis was performed several times in order to simulate the effect of sequencing depth. Results from all randomly generated datasets correlated with the original dataset (spearman’s rho ranged from 0.95 to 0.99). The variation around the line of the best fitting original dataset increased with smaller dataset sizes (Figure 7A) and thus, the mean squared differences decreased with dataset size (Supplementary file 9). The Kruskal-Wallis rank sum test shows significant differences between the different rarefied datasets regarding to the squared differences (χ2= 125.9, p-value <0.0001). Supplementary file 10 shows that the datasets with 200 and 500 random sequences were significantly different to the larger dataset sizes with 5000 and 7712 sequences (p-value ≤0.0034), whereas datasets with 1000, 5000 and 7712 showed no significant differences to each other (Supplementary file 10). The lowest average hit ratio of 53.6% was reached in the datasets with 200 random sequences and the highest average hit ratio of 89.8% was determined for the datasets with 7712 random sequences (Supplementary file 9; Figure 7B). A significant difference between all the datasets with regards to the hit ratio was identified ((χ2= 90.9, p-value <0.0001). The hit ratio for the dataset with 200 and 500 random sequences is significantly different to the larger dataset sizes with 5000 and 7712 sequences (p-value ≤0.0079). The average unknown classification rate of contamination decreased with increasing number of sequences used i.e. from 45.4% for the dataset with 200 sequences to 28.4% with 7712 sequences (Figure 7B). The average difference of unknown classified contamination compared to the original dataset ranged from 18.2% (dataset size 200) to 1.5% (dataset size 7712; Supplementary file 9). The Kruskal-Wallis test detected significant differences between the differences in the amount of unknown classified contaminations between the dataset sizes (χ2= 93.9, p-value <0.0001), which can be mainly assigned to the difference between datasets with 200 and 500 sequences vs. 5000 and 7712 sequences (Supplementary file 10).

**Figure 7:**
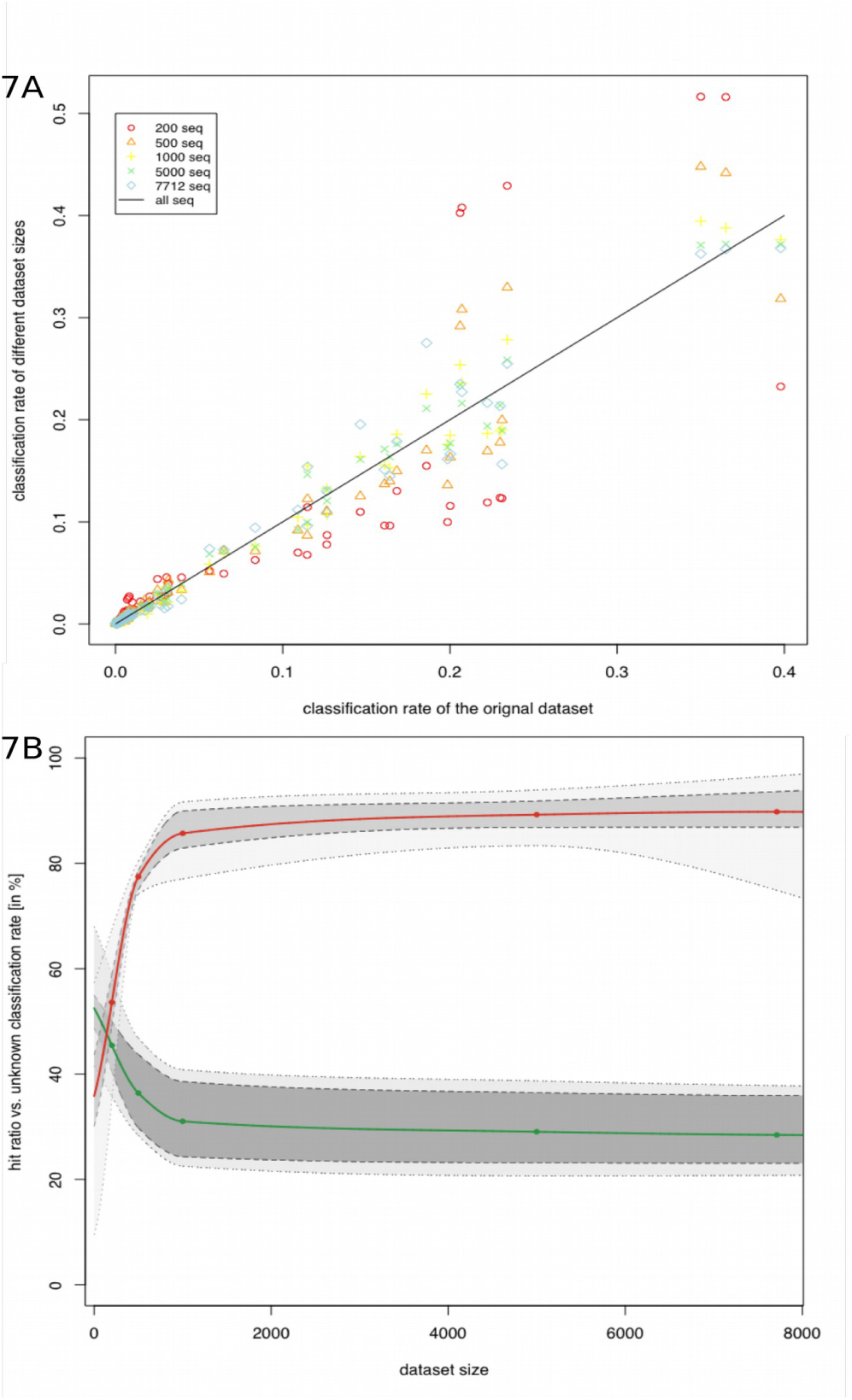
**(A):** The correlation of the different dataset sizes compared to original dataset. **(B):** The hit ratio (red points) and unknown classified sequences (green points) for the different used dataset sizes, with extrapolation values for not tested dataset sizes shown as solid lines. Dotted lines presented the min-max values and the dashed lines indicated the 25th and 75th percentile of-hit ratio and unknown classified sequences.

## Discussion

High throughput sequencing technologies have proven to be a powerful approach to explore microbial communities in a large variety of natural habitats as well as in built environments. The decreasing costs of these tools now also offer new perspectives to implement them in food production in order to investigate the impact of a given shift on the microbiota and their roles in a food system, which is directly correlated to food safety, food shelf life, flavour, and many other aspects (Bokulich et al. 2016). Here we present the first study to utilise the high-throughput, long-read sequencing capability of the PacBio technology to obtain thousands of full-length 16S rRNA gene sequences for microbial diversity studies during pig meat processing. Although PacBio sequencing is not as cost-effective as some of the available short-read platforms such as Illumina, it is able to produce longer read lengths, resulting in higher resolution for taxonomic classification and microbial source tracking.

First, we determined bacterial cell numbers with current standard techniques (Aerobic mesophilic and *Enterobacteriaceae* counts) and with additional methods (*Pseudomonadaceae* and BCE counts) to get a framework of the overall microbiological status of the slaughterhouse. This initial assessment revealed similar trends along the processing line as past investigations and showed high microbial numbers for several surface samples (Spescha et al. 2006; Warriner et al. 2002). The high bacterial numbers on the polishing tunnel equipment and the significant increase of AMC, and *Pseudomonadaceae* counts on the meat from singeing to after polishing already indicate transfer of bacteria at this step. A significant increase in bacterial contamination after polishing was also reported by Wheatley et al. (Wheatley et al. 2014). It is likely that the bacteria that survived the singeing treatment get spread over the carcass during polishing in addition of transfer of bacteria that persist in the polishing equipment (Gill 2005). Basic community analysis of the 16S rRNA gene sequencing data also exposed a higher microbial diversity after the polishing step when compared to after singeing, further indicating the same step as a possible transmission event.

When the pigs entered the facility, they had a high variation of the microbial community composition on the skin which points out large differences of the individual pig skin microbiota. However, this variation is greatly reduced after the singeing step. Thus, the initial microbial community structure on the carcass surface has little impact on the effectiveness of decontamination measures like singeing, resulting in the establishment of a similar community on each carcass. The community then remains relatively stable until the loading station where it starts to diversify again. This can be explained by the cooling period (16h at 7°C) between the classification and truck step. Small variations in the psychrotolerant part of the microbial community could lead to disparate alterations during the cooling period.

The 50 most abundant ASVs that were detected are associated to genera that have been previously found within the meat processing environment and some of them, e.g. *Bacillus, Chryseobacterium, Flavobacterium, Lactococcus, Microbacterium*, and *Moraxella* are considered to have spoilage potential (de Smidt 2016; Hultman et al. 2015; de Filippis et al. 2013; A. Davies and Board 1998). Interestingly, the meat samples were dominated by a number of ASVs associated to the genus *Anoxybacillus* after the singeing step. This genus consists of 22 species that were isolated from hot springs or manure, but were also regularly detected in dairy and meat processing environments (Khan et al. 2018; Pikuta et al. 2000; Burgess et al. 2010; Hultman et al. 2015). *Anoxybacillus* are aerobic or facultative anaerobic spore-formers, with an optimum growth at 50–65°C and neutral pH, are alkalitolerant, and are able to form biofilms (Goh et al. 2013; Burgess et al. 2009). Thus, they are able to survive the heat treatment (Scalding and singeing), explaining their high relative abundance on the meat after the singeing step. Overall, the microbial community on the meat samples (Singeing – Truck) was vastly different from the community on the skin of the animals when they entered the facility (Sticking). Hence, we hypothesized that the majority of the microorganisms were transferred onto the meat during processing and that only a smaller portion of the skin microbiota persists on the meat surface. Indeed, we concluded that more than half of the 500 most abundant ASVs were supposed to originate from water or soil habitats, rather than from animal-associated sources, suggesting that they were eventually transferred during processing.

We then used the software SourceTracker, which applies a Bayesian framework to estimate the proportion of each source contributing to a designated sink sample, to trace transmission routes and track down the sources of these microorganisms (Knights et al. 2011). This tool has been widely used to map microbial populations or gene flows in a variety of ecosystems, from coastal waters and drinking water systems to ATM keypads in New York, neonatal intensive care units, and to global antibiotic resistance gene pollution over diverse environmental types (Henry et al. 2016; Liu et al. 2018; Bik et al. 2016; Hewitt et al. 2013; Li et al. 2018). Surprisingly, only one study applied SourceTracker to a dataset obtained from a food environment (Brewery) so far (Bokulich et al. 2015). Here we present the first study using SourceTracker on full-length 16S rRNA gene sequencing data which is also the first applying it in the scope of a meat processing environment. Our analysis revealed key microbial transmission sites throughout the facility that were not identified with the current standard techniques. The main contamination sources contributing to the microbial community found on meat were the polishing tunnel equipment, gloves of employees, and a railing at the classification step. The polishing tunnel was identified as a critical operation during pork slaughtering in the past (Yu et al. 1999; Bolton et al. 2002). However, the gloves and railings are generally not considered as such and had low microbial levels indicating good hygiene practices. Still a lot of microorganisms were transferred from these positions. A closer look at the taxonomy of the transferred bacteria exposed unique transmission patterns for individual taxa. Noticeably, particular species occupy different environmental niches across the facility showing the importance of high taxonomic resolution for microbial source tracking in food processing plants. In fact, whole genome sequencing or metagenomic shotgun sequencing have been proposed to achieve strain-level resolution and are thought to be necessary to track microbes during food processing (Jagadeesan et al. 2019; Bergholz et al. 2014; Leonard et al. 2015). While we were not able to identify potential pathogens or specific spoilage organisms on a strain level, our approach displays that full-length 16S rRNA gene sequencing delivers a deep enough resolution for environmental monitoring within a facility, since we were able to distinguish between different microbial populations that were transferred from respective sources. Thus, strain-level tracking using metagenomics is still essential for molecular epidemiology and diagnostics but it might be overkill for monitoring systems (Allard et al. 2018).

Moreover, we were able to show that it is not necessary to deeply sequence the amplicon libraries, but that datasets with 1000 sequences/sample provide a comparable result to deeper sequenced datasets. Thus it is feasible to multiplex many samples on a single sequencing run substantially decreasing costs, which essentially accomplishes affordability for regular monitoring checks. However, several challenges and limitations remain before we can realize the full potential of NGS techniques as food safety applications. Currently, bioinformatics workflows are implemented and executed based on ad hoc lab specific experiences. Each lab uses different protocols and the documentation of the actual process is rarely well recorded or presented. Hence, it is paramount to expose, formalize, and standardize sampling techniques, as well as workflows for bioinformatics pipelines, processing, and data management (Weimer et al. 2016). In that way, it would be possible to compare datasets and leverage systematic authentication of the microbiome and its variation throughout the supply chain to understand microbial contamination during food production on a broader scale.

## Conclusion

Since we observed only one significant increase in microbial numbers along the slaughter line, we consider AMC as good general hygiene indicators that can reflect substantial contamination incidents, but they fail in describing more complex population flows. Thus, AMC and EB determination by microbiological reference methods is not sufficient to detect the full extent of microbial transmission events. This study showcases that high-throughput full-length 16S rRNA gene sequencing can reveal valuable information about the microbial communities in pork production plants and expose critical contamination steps during slaughtering. We were able to pinpoint many taxa to specific sources, facilitating targeted combat of potential pathogens or meat-spoilage organisms in the analysed facility. Our findings contribute to improve or optimise hygiene standards in the meat industry to further minimize the risk of microbial cross-contamination. Furthermore, the methods used in this study can be applied to any other food-related industry to universally promote our knowledge about microbial transfer during food processing. Continuing advances in long-read sequencing technologies like the release of the Sequel II system and the development of full rRNA operon sequencing strategies will further increase the throughput and taxonomic resolution, offering great potential to implement them in monitoring systems (Martijn et al. 2019).

## Supporting information

Detailed list of samples taken at the slaughterhouse and type of analysis performed

Details on the PCR setup for the 16S rRNA gene qPCR

R markdown script of the sequence processing and community analysis workflow

Rarefaction curve of individual 16S rRNA gene libraries

Prevalence of genera comprising top 50 ASVs

Relative abundances of all detected genera across all meat positions

Relative abundances of all detected genera across all surface positions

Phylogenetic tree of 16S rRNA gene sequences associated to the genus Chryseobacterium

Mean squared differences, mean hit ratios, mean unknown classification rates and mean differences of unknown classification rates

Results of the Dunn tests after a significant Kruskal-Wallis rank sum test for mean squared differences, hit ratio and unknown classification rate

## Acknowledgements

We thank Viktoria Neubauer for fruitful discussions and proofreading of the manuscript and Nikolaus Pfisterer and Christian Mattes for their assistance during sampling. Additionally, we thank Lauren Alteio for R package recommendations and long-distance emotional support.

## Authors’ contributions

BZ, SW, MW, and EM conceived and designed the study. Sampling was performed by BZ, SW, IR, and BS. BZ, IR, ST, MD, and BS performed the experiments and BZ, ED, CS, and AZ developed bioinformatics pipelines. Data analysis and statistics were performed by BZ, EM, BP, and FFR, and BZ and EM wrote the manuscript. All authors read and approved the final manuscript.

## Supplementary file legends

Supplementary file 1: D*etailed list of samples taken at the slaughterhouse and type of analysis performed*

Supplementary file 2: *Details on the PCR setup for the 16S rRNA gene qPCR*

Supplementary file 3: *R script of the sequence processing and community analysis workflow*

Supplementary file 4: *Rarefaction curve of individual 16S rRNA gene libraries*

Supplementary file 5: *Prevalence of genera comprising top 50 ASVs*

Supplementary file 6: *Relative abundances of all detected genera across all meat positions*

Supplementary file 7: *Relative abundances of all detected genera across all surface positions*

Supplementary file 8: *Phylogenetic tree of 16S rRNA gene sequences associated to the genus Chryseobacterium. The size of the circles at the tips of the tree indicate the relative abundance at the respective position illustrated by the colours.*

Supplementary file 9: *Mean squared differences, mean hit ratios, mean unknown classification rates and mean differences of unknown classification rates compared to the original dataset, stratified by the different dataset size. N.B. All presented means in the table were calculated as the difference between the randomly generated datasets and the original dataset.*

Supplementary file 10: *Results of the Dunn tests after a significant Kruskal-Wallis rank sum test for mean squared differences, hit ratio and unknown classification rate, whereby the original dataset was used as reference. Significant p-values are highlighted in bold.*

