## Supplementary figures and images for "The sources and transmission routes of microbial populations throughout a meat processing facility"

### Phylogenetic tree of 16S rRNA gene sequences associated to the genus Chryseobacterium

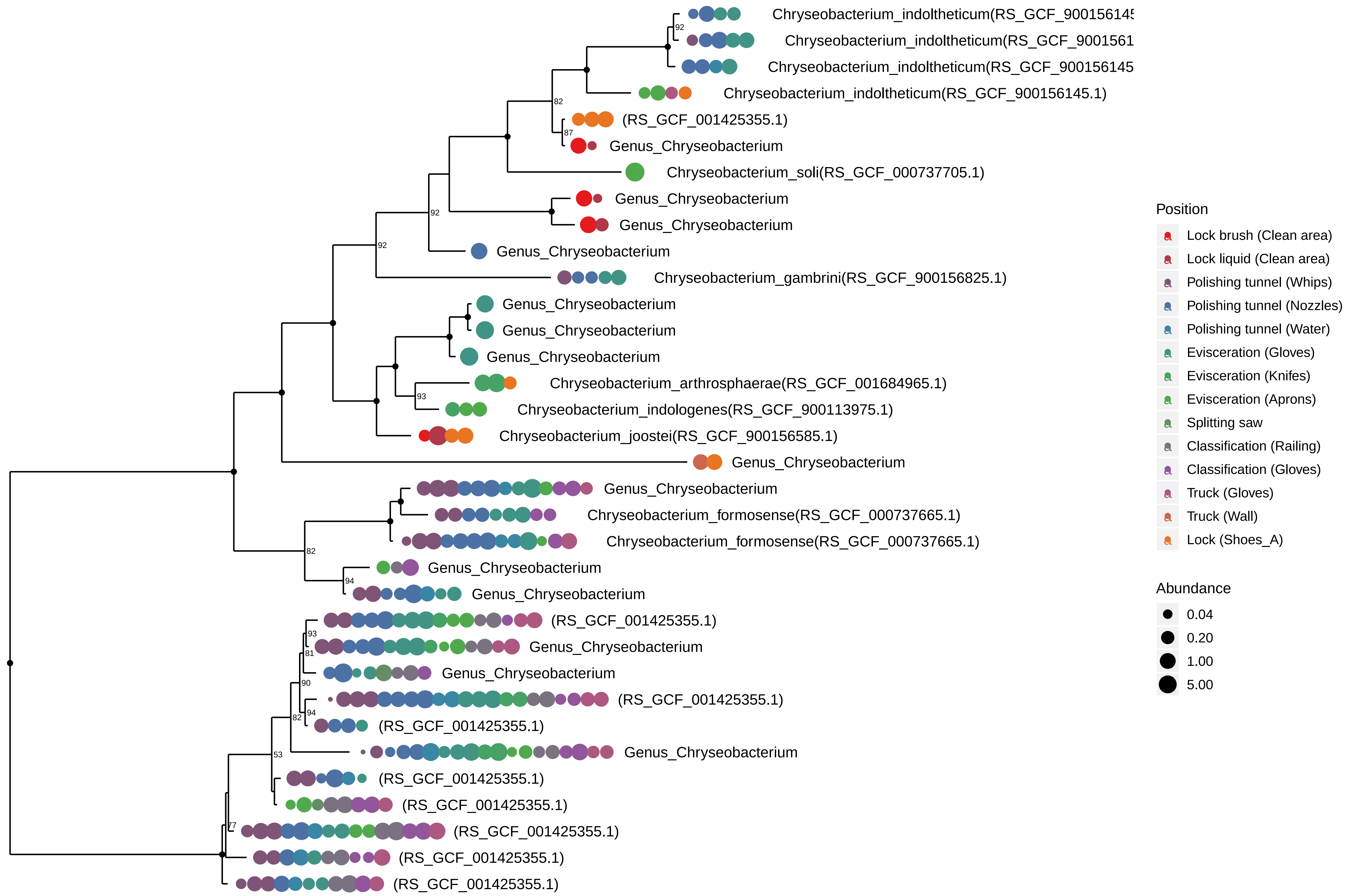

### Rarefaction curve of individual 16S rRNA gene libraries

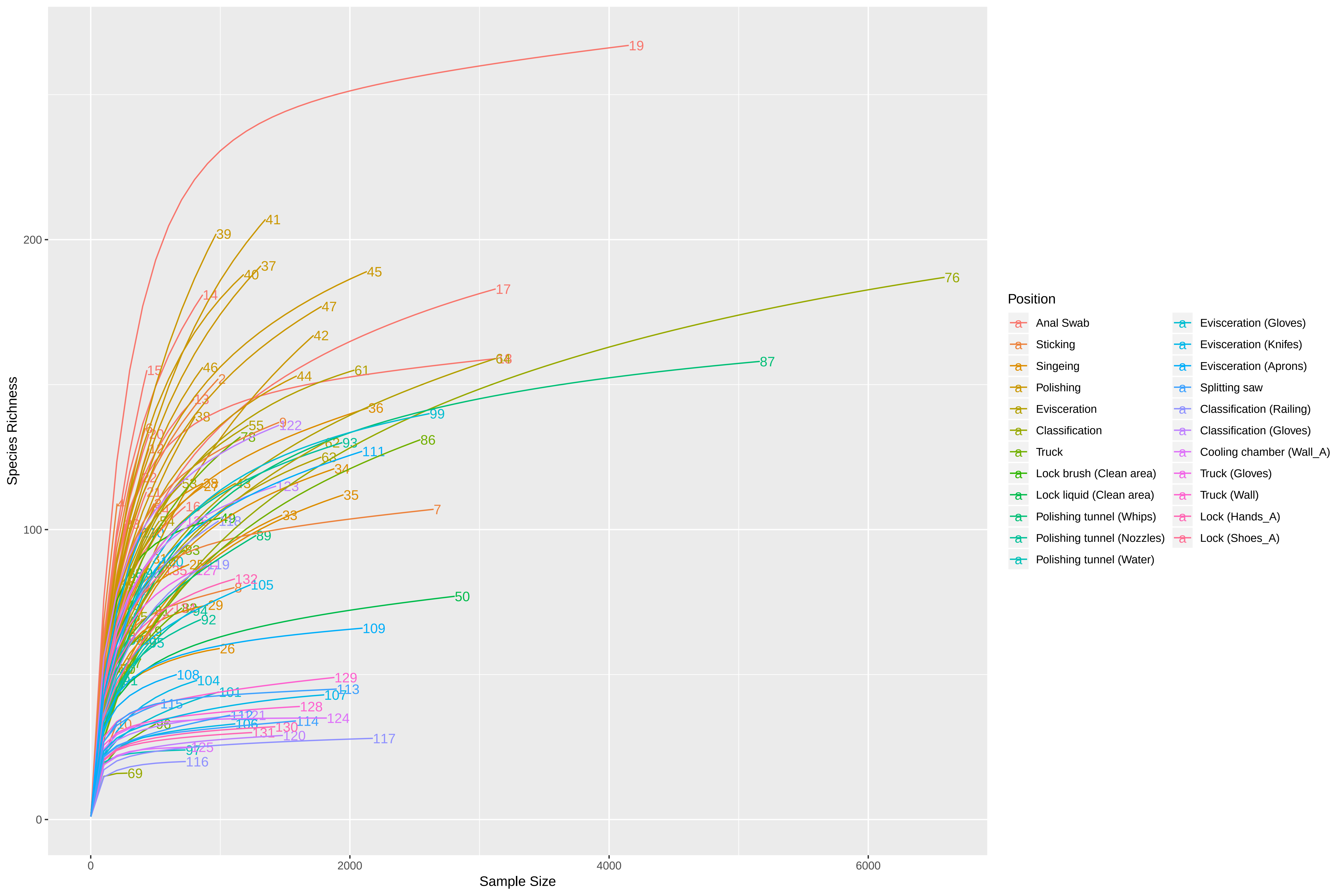
