## Supplementary material for "The sources and transmission routes of microbial populations throughout a meat processing facility": Relative abundances of all detected genera across all meat positions

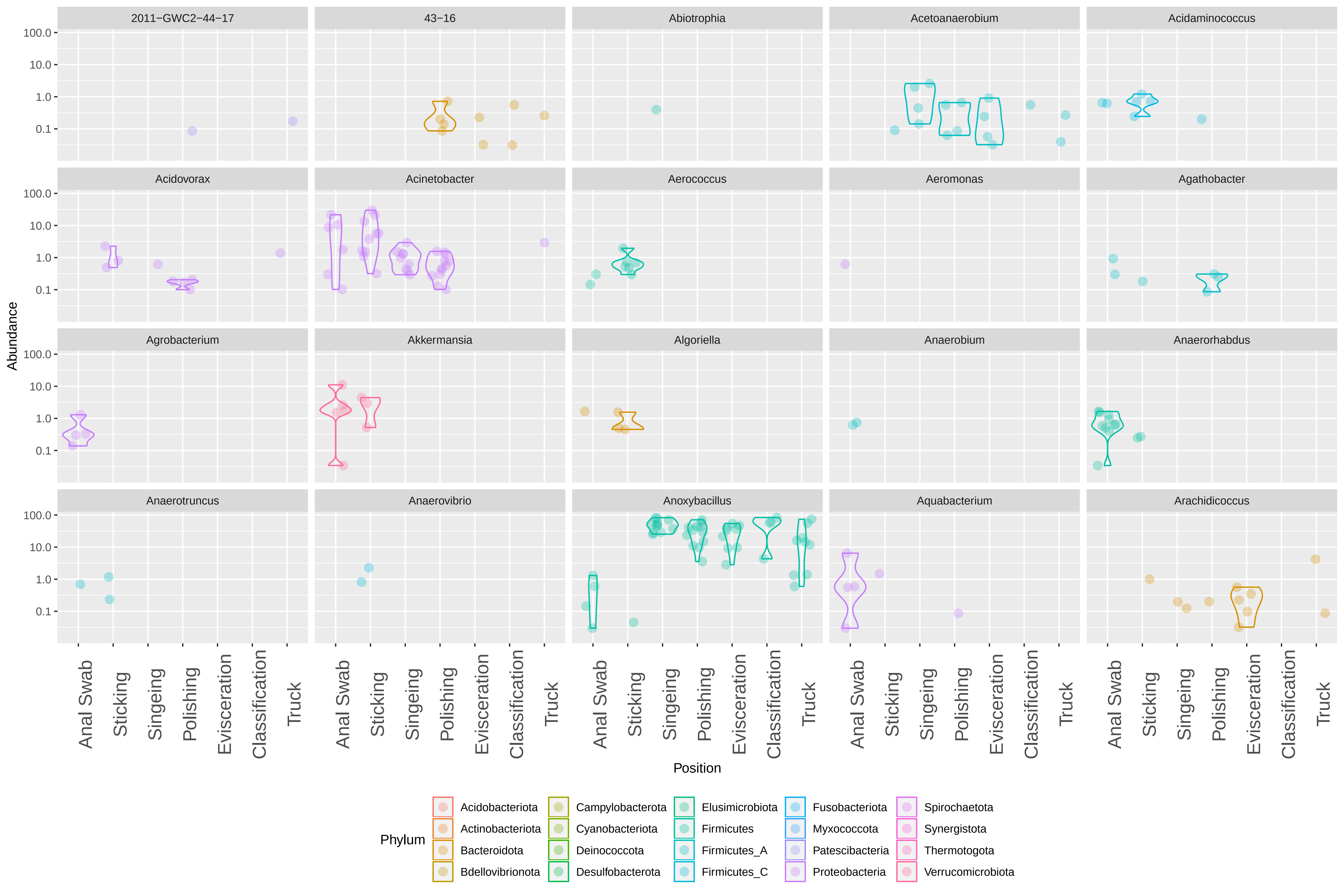
2011−GWC2−44−17
43−16
Abiotrophia
Acetoanaerobium
Acidaminococcus
Acidovorax
Acinetobacter
Aerococcus
Aeromonas
Agathobacter
Agrobacterium
Akkermansia
Algoriella
Anaerobium
Anaerorhabdus
Anaerotruncus
Anaerovibrio
Anoxybacillus
Aquabacterium
Arachidicoccus
Abundance
100.0
10.0
1.0
0.1
Anal Swab
Sticking
Singeing
Polishing
Evisceration
Classification
Truck
Position
Phylum
Acidobacteriota
Actinobacteriota
Bacteroidota
Bdellovibrionota
Campylobacterota
Cyanobacteriota
Deinococcota
Desulfobacterota
Elusimicrobiota
Firmicutes
Firmicutes_A
Firmicutes_C
Fusobacteriota
Myxococcota
Patescibacteria
Proteobacteria
Spirochaetota
Synergistota
Thermotogota
Verrucomicrobiota

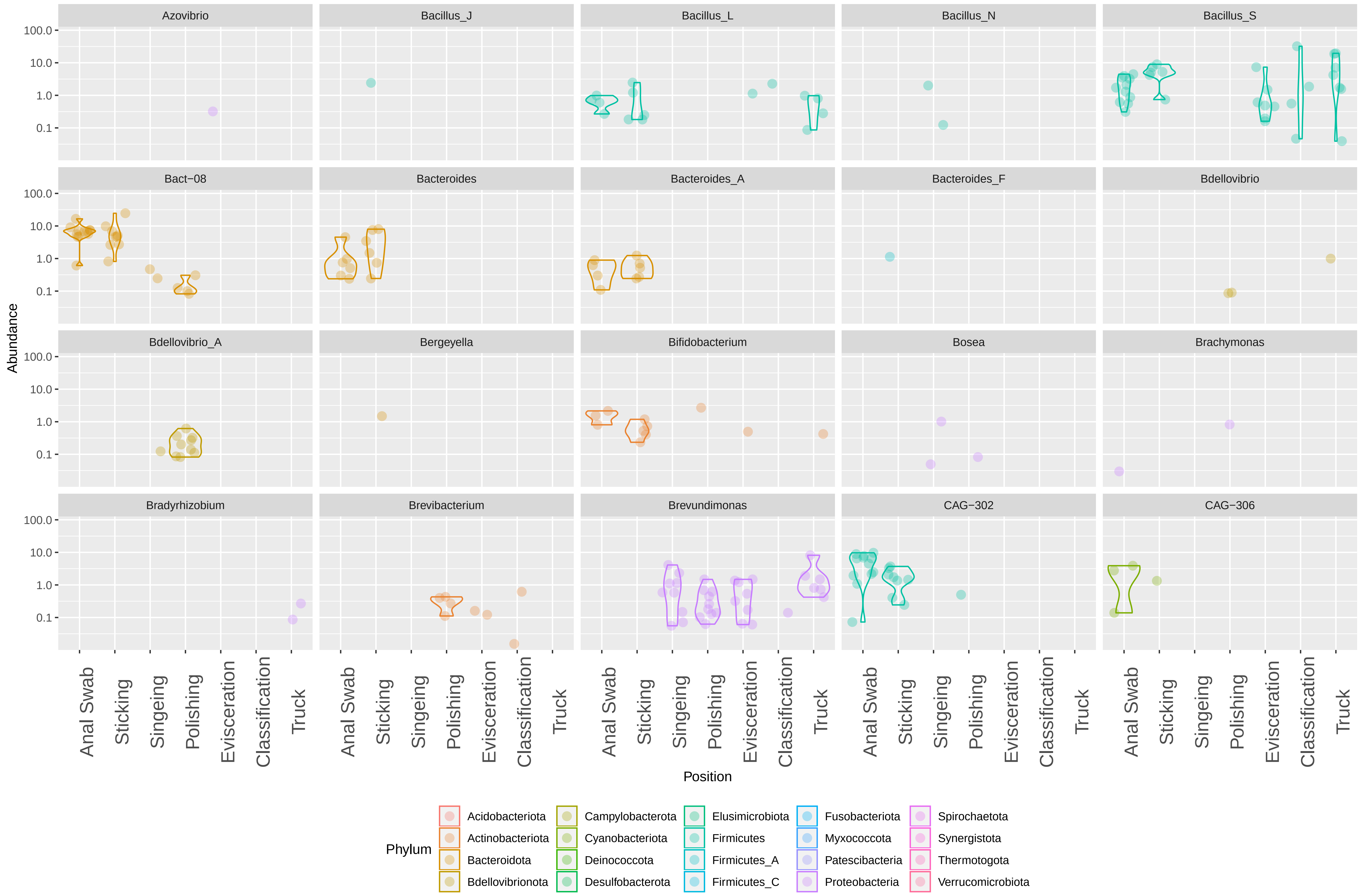
Azovibrio
Bacillus_J
Bacillus_L
Bacillus_N
Bacillus_S
Bact−08
Bacteroides
Bacteroides_A
Bacteroides_F
Bdellovibrio
Bdellovibrio_A
Bergeyella
Bifidobacterium
Bosea
Brachymonas
Bradyrhizobium
Brevibacterium
Brevundimonas
CAG−302
CAG−306
Abundance
100.0
10.0
1.0
0.1
Anal Swab
Sticking
Singeing
Polishing
Evisceration
Classification
Truck
Position
Phylum
Acidobacteriota
Actinobacteriota
Bacteroidota
Bdellovibrionota
Campylobacterota
Cyanobacteriota
Deinococcota
Desulfobacterota
Elusimicrobiota
Firmicutes
Firmicutes_A
Firmicutes_C
Fusobacteriota
Myxococcota
Patescibacteria
Proteobacteria
Spirochaetota
Synergistota
Thermotogota
Verrucomicrobiota

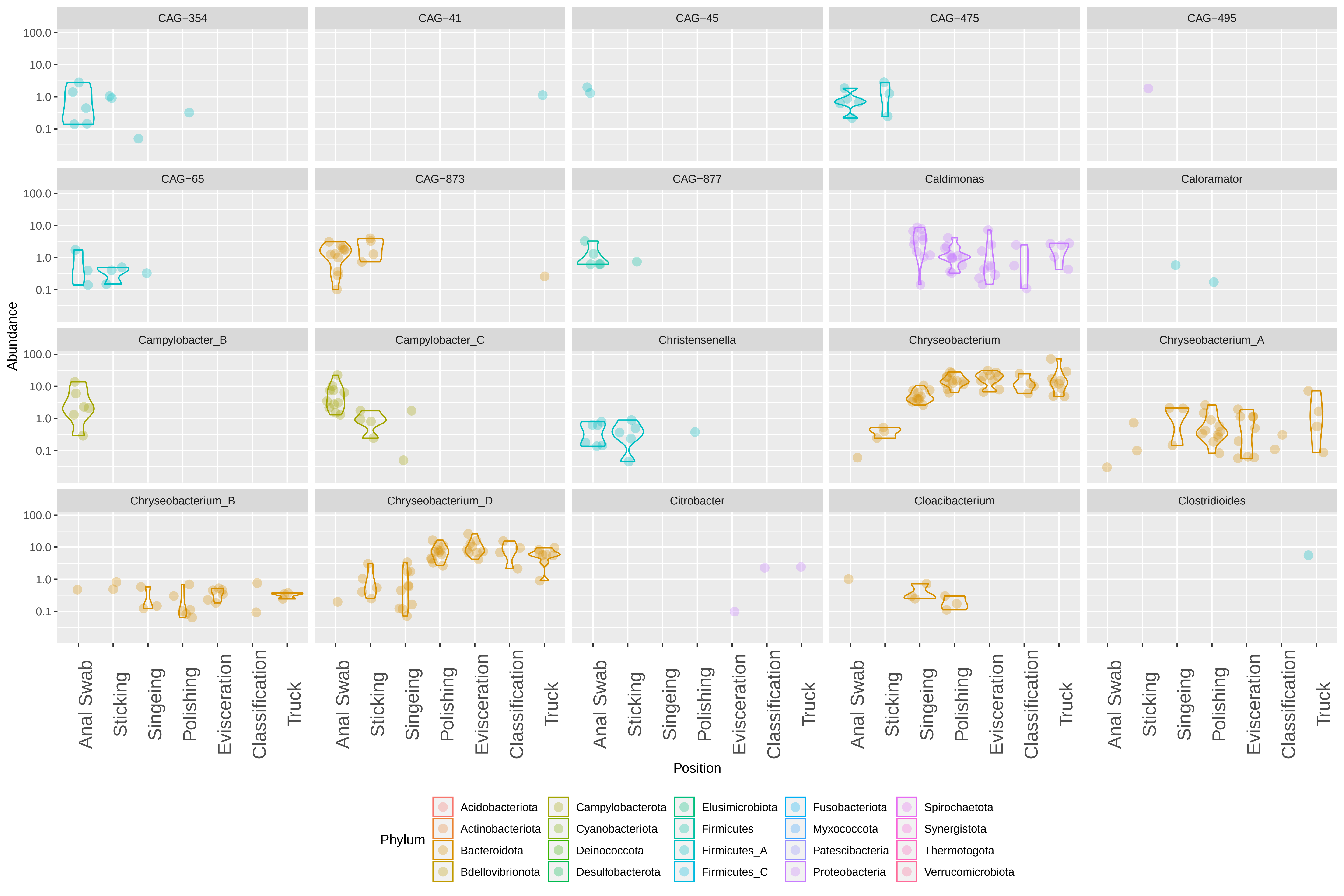
CAG−354
CAG−41
CAG−45
CAG−475
CAG−495
CAG−65
CAG−873
CAG−877
Caldimonas
Caloramator
Campylobacter_B
Campylobacter_C
Christensenella
Chryseobacterium
Chryseobacterium_A
Chryseobacterium_B
Chryseobacterium_D
Citrobacter
Cloacibacterium
Clostridioides
100.0
10.0
1.0
0.1
Abundance
Anal Swab
Sticking
Singeing
Polishing
Evisceration
Classification
Truck
Position
Phylum
Acidobacteriota
Actinobacteriota
Bacteroidota
Bdellovibrionota
Campylobacterota
Cyanobacteriota
Deinococcota
Desulfobacterota
Elusimicrobiota
Firmicutes
Firmicutes_A
Firmicutes_C
Fusobacteriota
Myxococcota
Patescibacteria
Proteobacteria
Spirochaetota
Synergistota
Thermotogota
Verrucomicrobiota

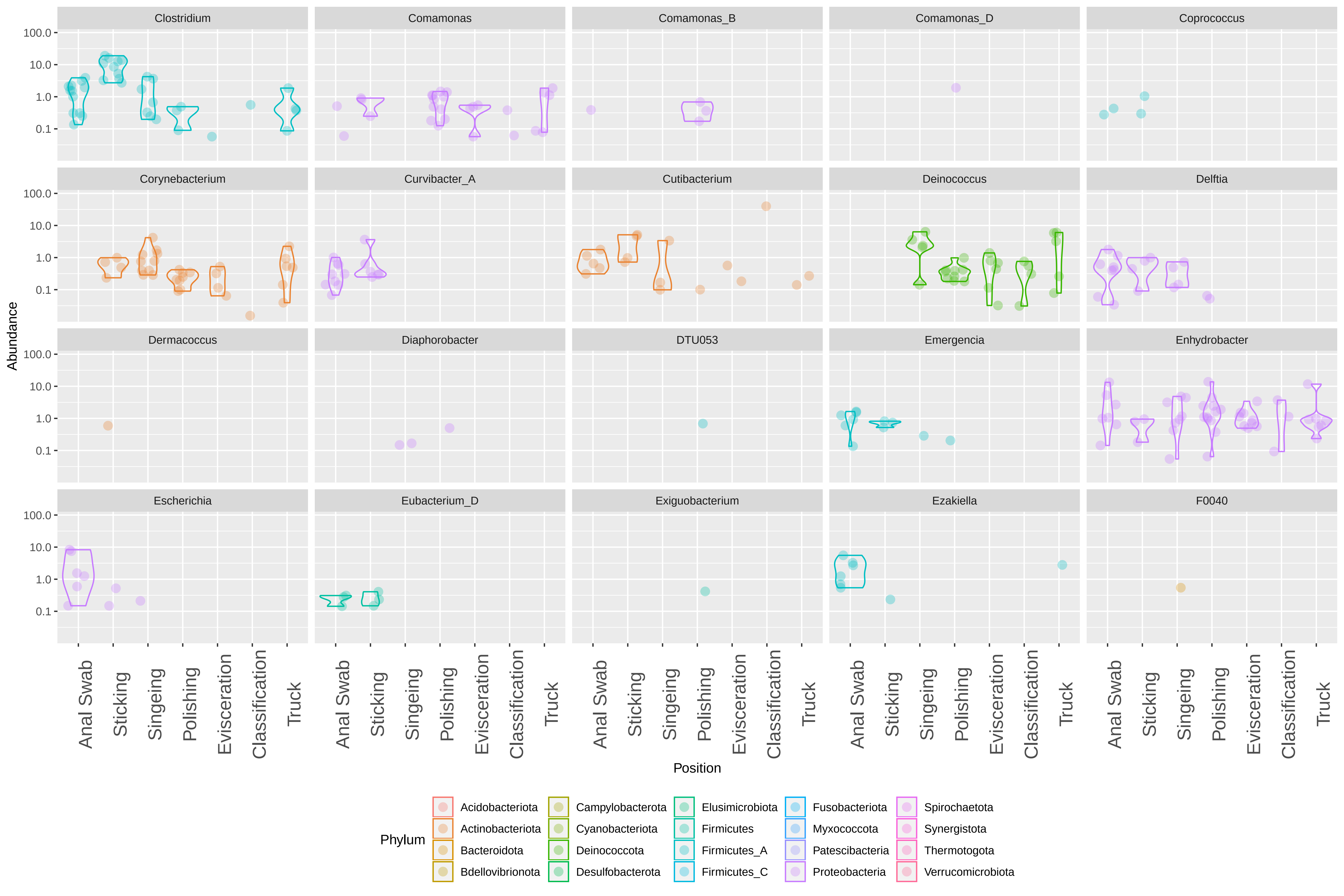
Clostridium
Comamonas
Comamonas_B
Comamonas_D
Coprococcus
Corynebacterium
Curvibacter_A
Cutibacterium
Deinococcus
Delftia
Dermacoccus
Diaphorobacter
DTU053
Emergencia
Enhydrobacter
Escherichia
Eubacterium_D
Exiguobacterium
Ezakiella
F0040
Abundance
100.0
10.0
1.0
0.1
Anal Swab
Sticking
Singeing
Polishing
Evisceration
Classification
Truck
Position
Phylum
Acidobacteriota
Actinobacteriota
Bacteroidota
Bdellovibrionota
Campylobacterota
Cyanobacteriota
Deinococcota
Desulfobacterota
Elusimicrobiota
Firmicutes
Firmicutes_A
Firmicutes_C
Fusobacteriota
Myxococcota
Patescibacteria
Proteobacteria
Spirochaetota
Synergistota
Thermotogota
Verrucomicrobiota

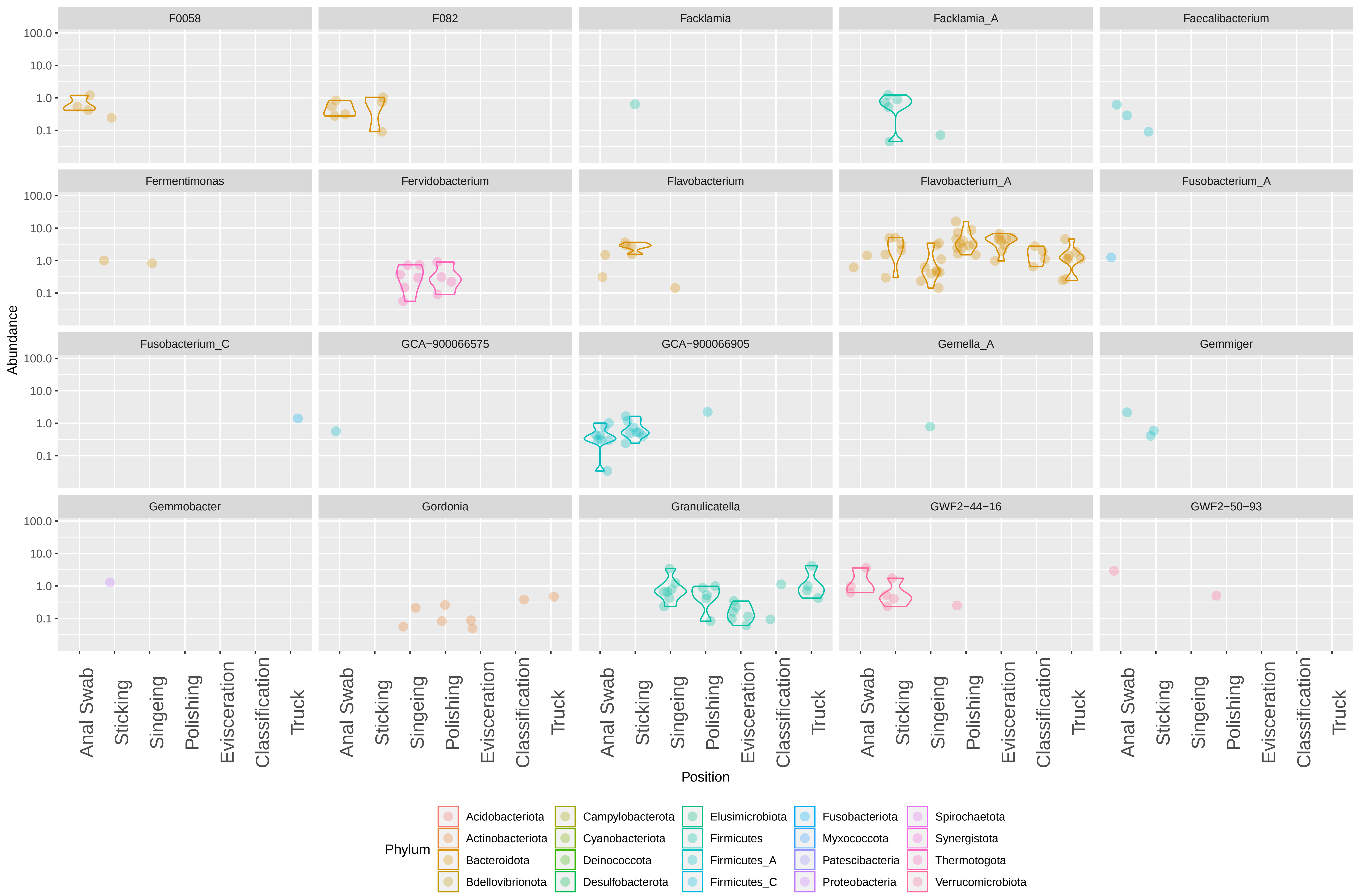

F0058
F082
Facklamia
Facklamia_A
Faecalibacterium
Fermentimonas
Fervidobacterium
Flavobacterium
Flavobacterium_A
Fusobacterium_A
Fusobacterium_C
GCA−900066575
GCA−900066905
Gemella_A
Gemmiger
Gemmobacter
Gordonia
Granulicatella
GWF2−44−16
GWF2−50−93
Abundance
100.0
10.0
1.0
0.1
Anal Swab
Sticking
Singeing
Polishing
Evisceration
Classification
Truck
Position
Phylum
Acidobacteriota
Actinobacteriota
Bacteroidota
Bdellovibrionota
Campylobacterota
Cyanobacteriota
Deinococcota
Desulfobacterota
Elusimicrobiota
Firmicutes
Firmicutes_A
Firmicutes_C
Fusobacteriota
Myxococcota
Patescibacteria
Proteobacteria
Spirochaetota
Synergistota
Thermotogota
Verrucomicrobiota

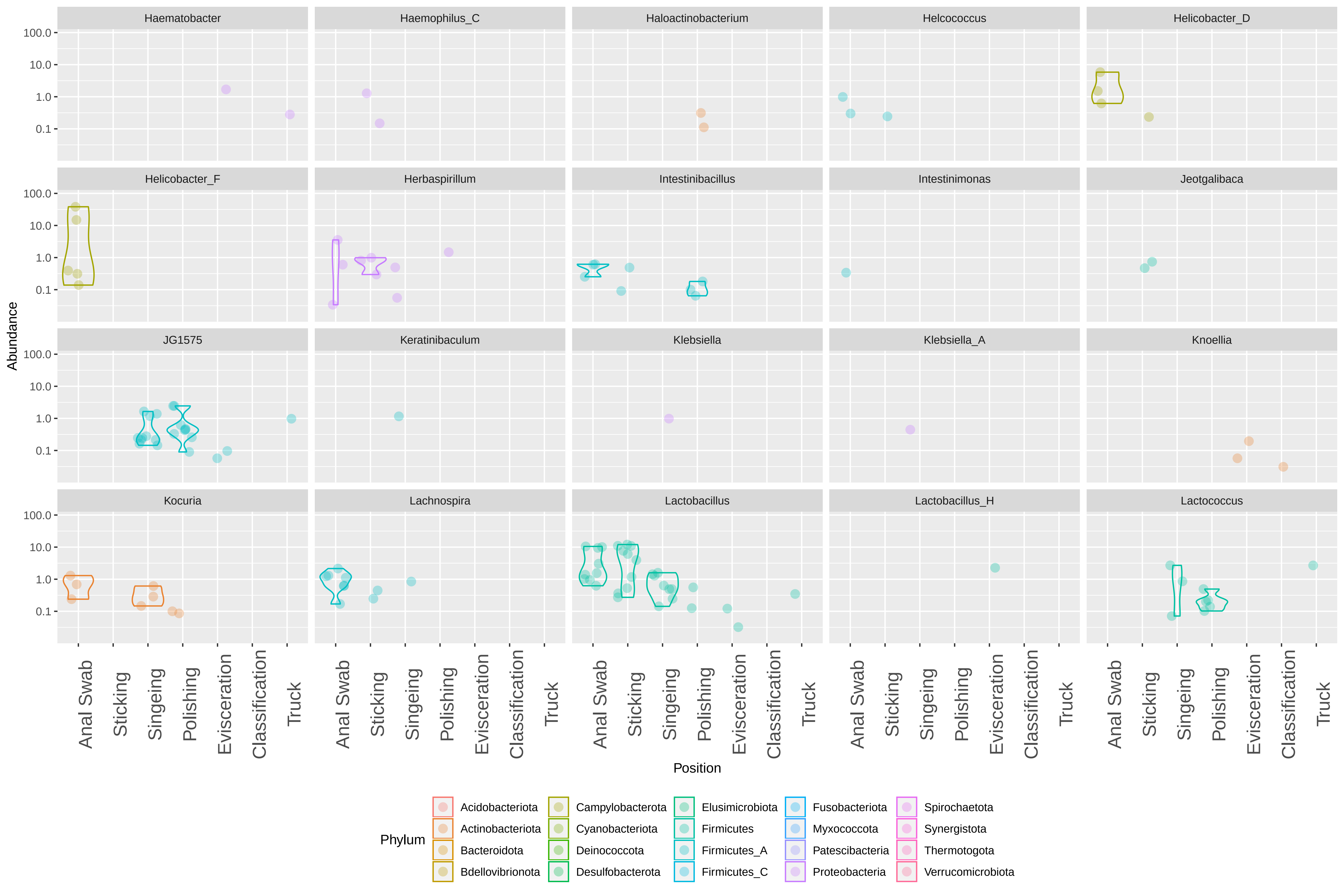

Haematobacter
Haemophilus_C
Haloactinobacterium
Helcococcus
Helicobacter_D
Helicobacter_F
Herbaspirillum
Intestinibacillus
Intestinimonas
Jeotgalibaca
JG1575
Keratinibaculum
Klebsiella
Klebsiella_A
Knoellia
Kocuria
Lachnospira
Lactobacillus
Lactobacillus_H
Lactococcus
Abundance
100.0
10.0
1.0
0.1
Anal Swab
Sticking
Singeing
Polishing
Evisceration
Classification
Truck
Position
Phylum
Acidobacteriota
Actinobacteriota
Bacteroidota
Bdellovibrionota
Campylobacterota
Cyanobacteriota
Deinococcota
Desulfobacterota
Elusimicrobiota
Firmicutes
Firmicutes_A
Firmicutes_C
Fusobacteriota
Myxococcota
Patescibacteria
Proteobacteria
Spirochaetota
Synergistota
Thermotogota
Verrucomicrobiota

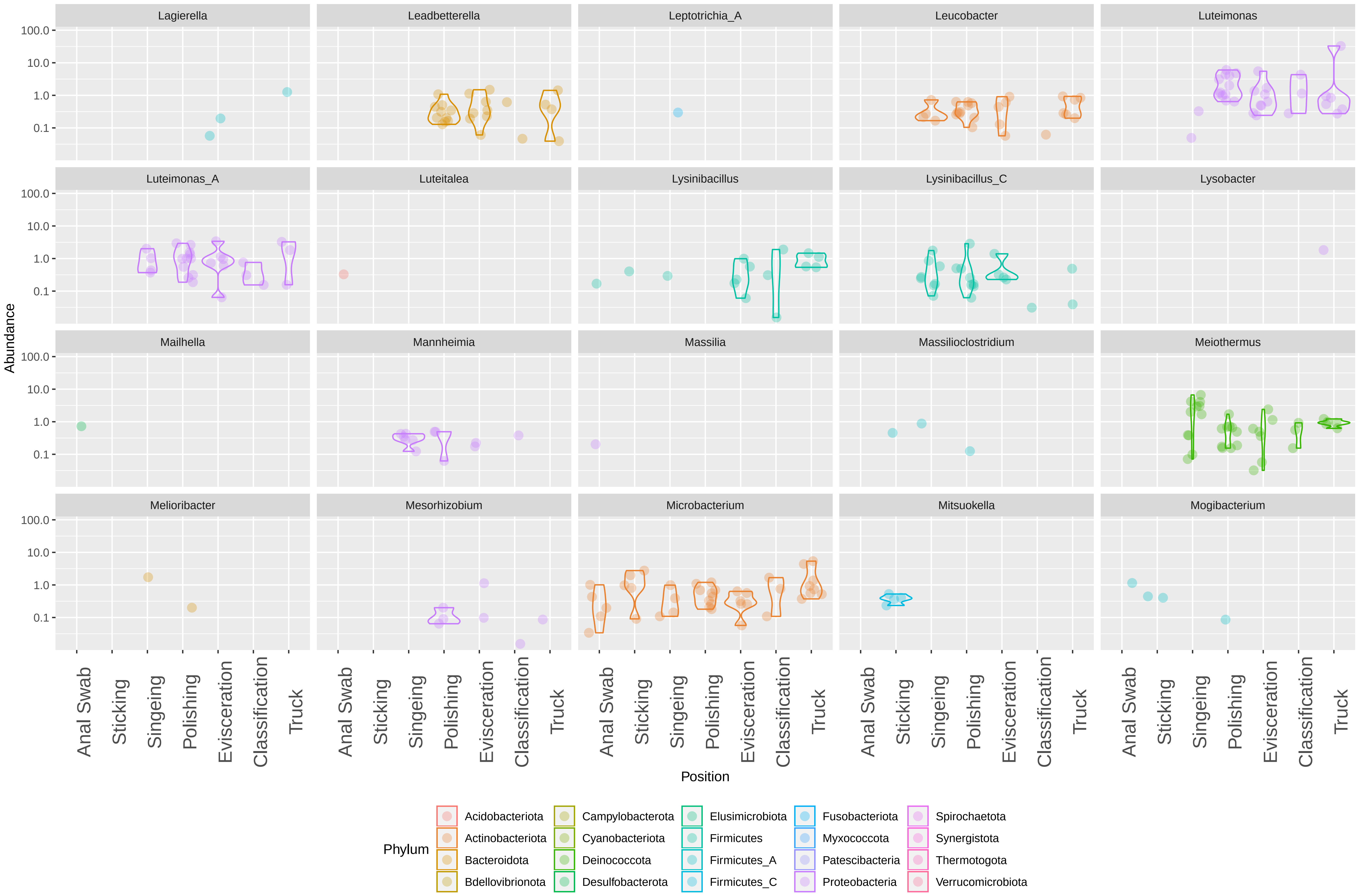

Lagierella
Leadbetterella
Leptotrichia_A
Leucobacter
Luteimonas
Luteimonas_A
Luteitalea
Lysinibacillus
Lysinibacillus_C
Lysobacter
Mailhella
Mannheimia
Massilia
Massilioclostridium
Meiothermus
Melioribacter
Mesorhizobium
Microbacterium
Mitsuokella
Mogibacterium
Abundance
100.0
10.0
1.0
0.1
Anal Swab
Sticking
Singeing
Polishing
Evisceration
Classification
Truck
Position
Phylum
Acidobacteriota
Actinobacteriota
Bacteroidota
Bdellovibrionota
Campylobacterota
Cyanobacteriota
Deinococcota
Desulfobacterota
Elusimicrobiota
Firmicutes
Firmicutes_A
Firmicutes_C
Fusobacteriota
Myxococcota
Patescibacteria
Proteobacteria
Spirochaetota
Synergistota
Thermotogota
Verrucomicrobiota

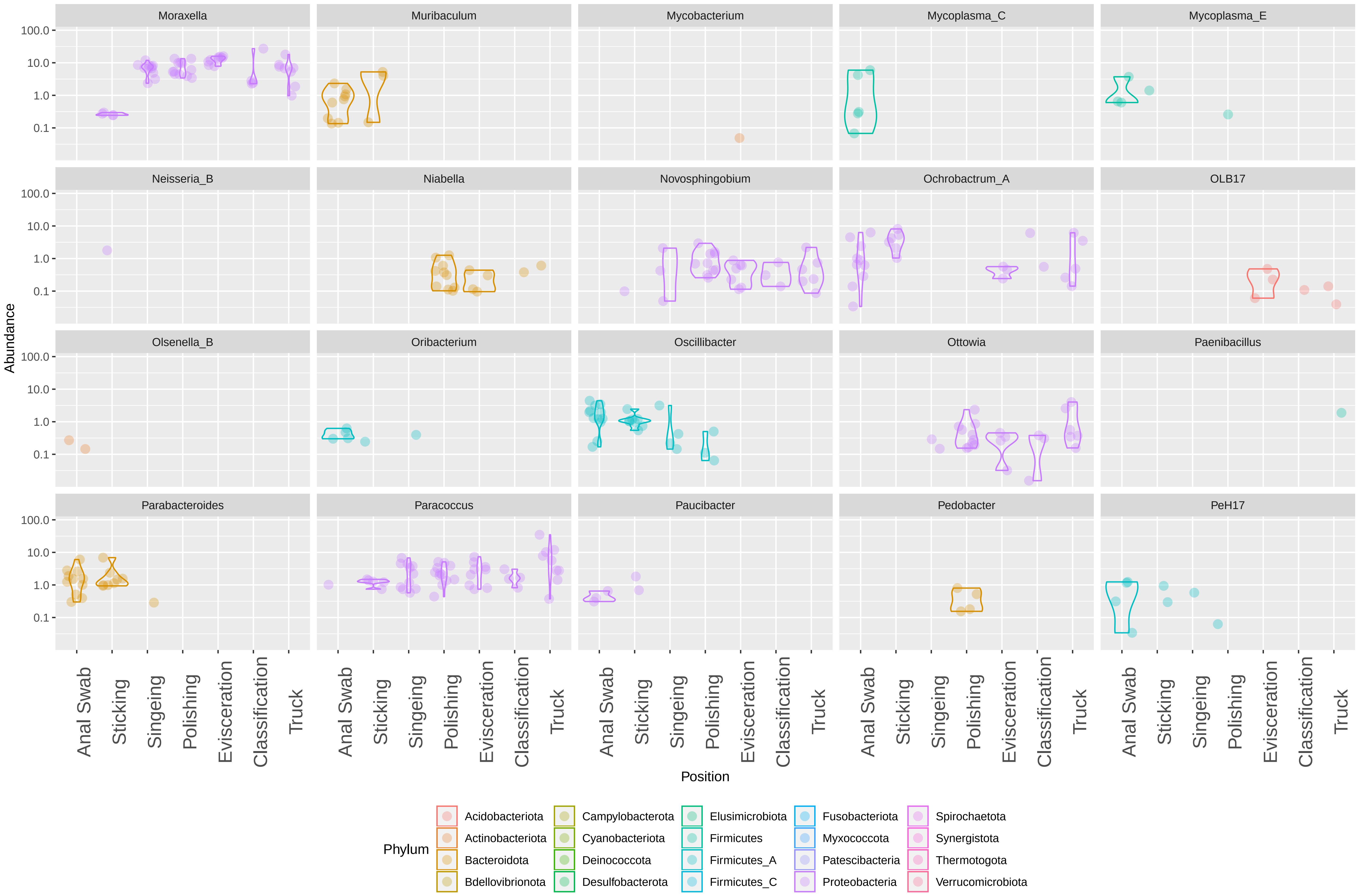

Moraxella
Muribaculum
Mycobacterium
Mycoplasma_C
Mycoplasma_E
Neisseria_B
Niabella
Novosphingobium
Ochrobactrum_A
OLB17
Olsenella_B
Oribacterium
Oscillibacter
Ottowia
Paenibacillus
Parabacteroides
Paracoccus
Paucibacter
Pedobacter
PeH17
Abundance
100.0
10.0
1.0
0.1
Anal Swab
Sticking
Singeing
Polishing
Evisceration
Classification
Truck
Position
Phylum
Acidobacteriota
Actinobacteriota
Bacteroidota
Bdellovibrionota
Campylobacterota
Cyanobacteriota
Deinococcota
Desulfobacterota
Elusimicrobiota
Firmicutes
Firmicutes_A
Firmicutes_C
Fusobacteriota
Myxococcota
Patescibacteria
Proteobacteria
Spirochaetota
Synergistota
Thermotogota
Verrucomicrobiota

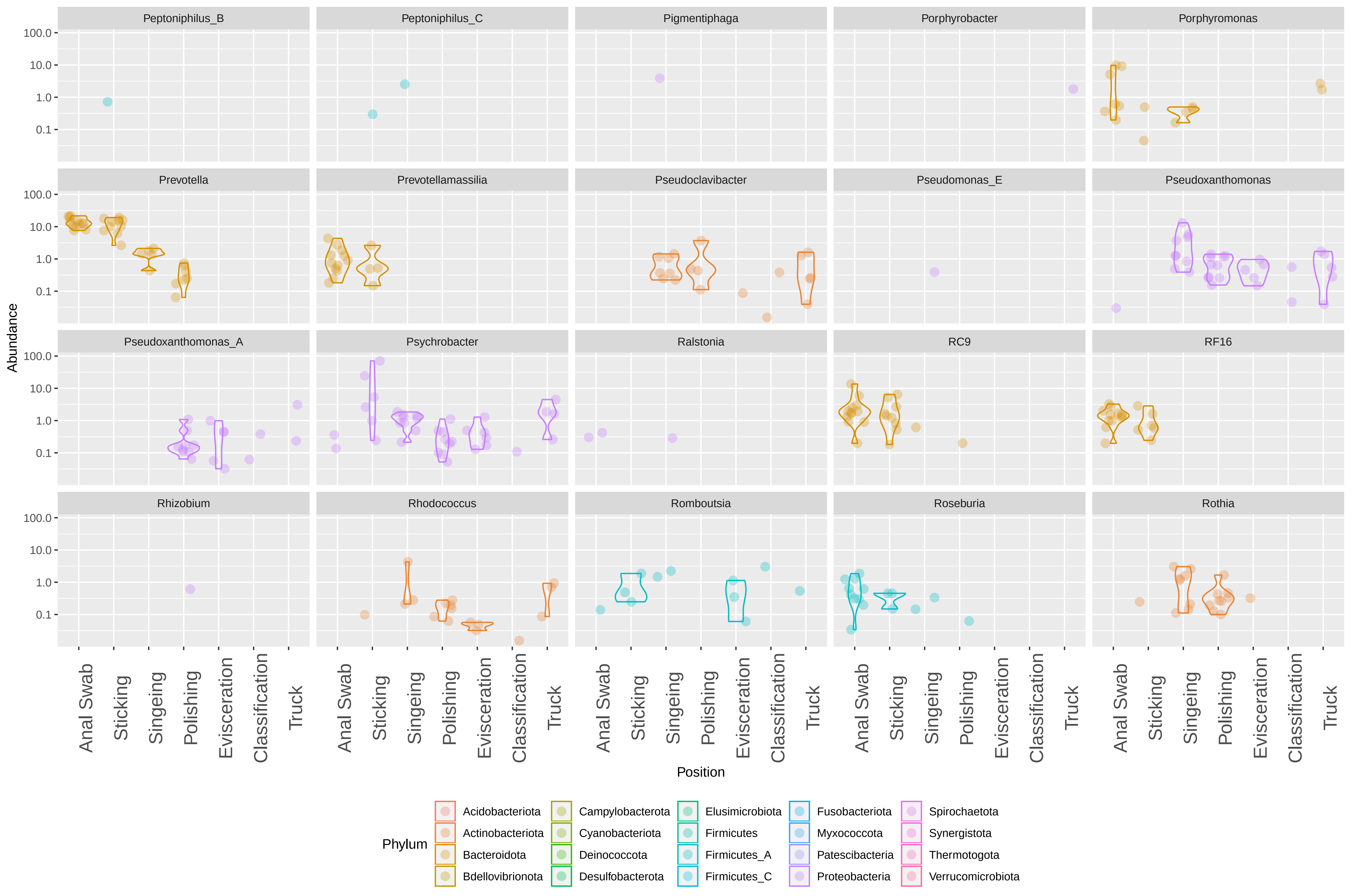
Peptoniphilus_B
Peptoniphilus_C
Pigmentiphaga
Porphyrobacter
Porphyromonas
Prevotella
Prevotellamassilia
Pseudoclavibacter
Pseudomonas_E
Pseudoxanthomonas
Pseudoxanthomonas_A
Psychrobacter
Ralstonia
RC9
RF16
Rhizobium
Rhodococcus
Romboutsia
Roseburia
Rothia
Abundance
100.0
10.0
1.0
0.1
Anal Swab
Sticking
Singeing
Polishing
Evisceration
Classification
Truck
Position
Phylum
Acidobacteriota
Actinobacteriota
Bacteroidota
Bdellovibrionota
Campylobacterota
Cyanobacteriota
Deinococcota
Desulfobacterota
Elusimicrobiota
Firmicutes
Firmicutes_A
Firmicutes_C
Fusobacteriota
Myxococcota
Patescibacteria
Proteobacteria
Spirochaetota
Synergistota
Thermotogota
Verrucomicrobiota

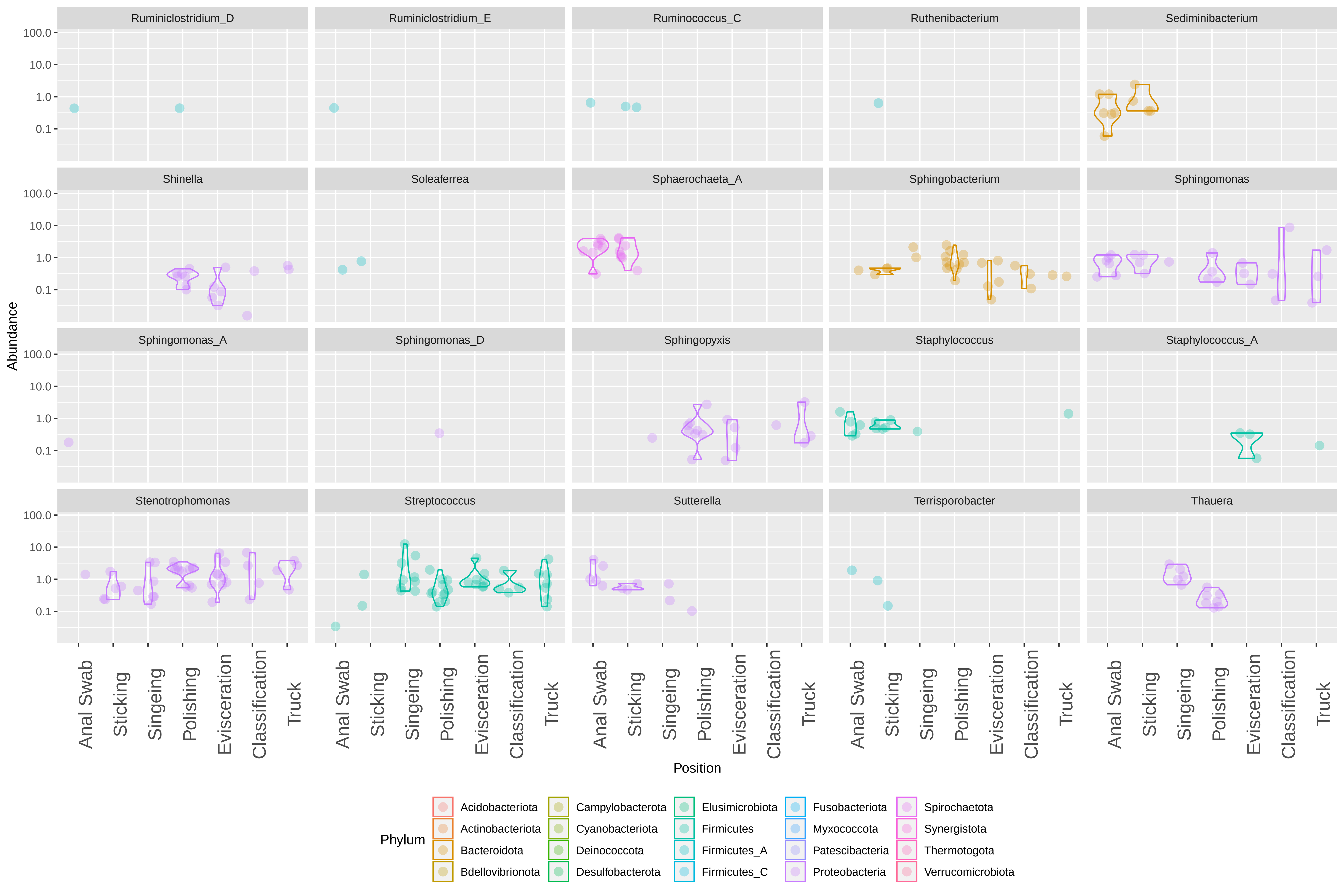
Ruminiclostridium_D
Ruminiclostridium_E
Ruminococcus_C
Ruthenibacterium
Sediminibacterium
Shinella
Soleaferrea
Sphaerochaeta_A
Sphingobacterium
Sphingomonas
Sphingomonas_A
Sphingomonas_D
Sphingopyxis
Staphylococcus
Staphylococcus_A
Stenotrophomonas
Streptococcus
Sutterella
Terrisporobacter
Thauera
100.0
10.0
1.0
0.1
Abundance
Anal Swab
Sticking
Singeing
Polishing
Evisceration
Classification
Truck
Position
Phylum
Acidobacteriota
Actinobacteriota
Bacteroidota
Bdellovibrionota
Campylobacterota
Cyanobacteriota
Deinococcota
Desulfobacterota
Elusimicrobiota
Firmicutes
Firmicutes_A
Firmicutes_C
Fusobacteriota
Myxococcota
Patescibacteria
Proteobacteria
Spirochaetota
Synergistota
Thermotogota
Verrucomicrobiota

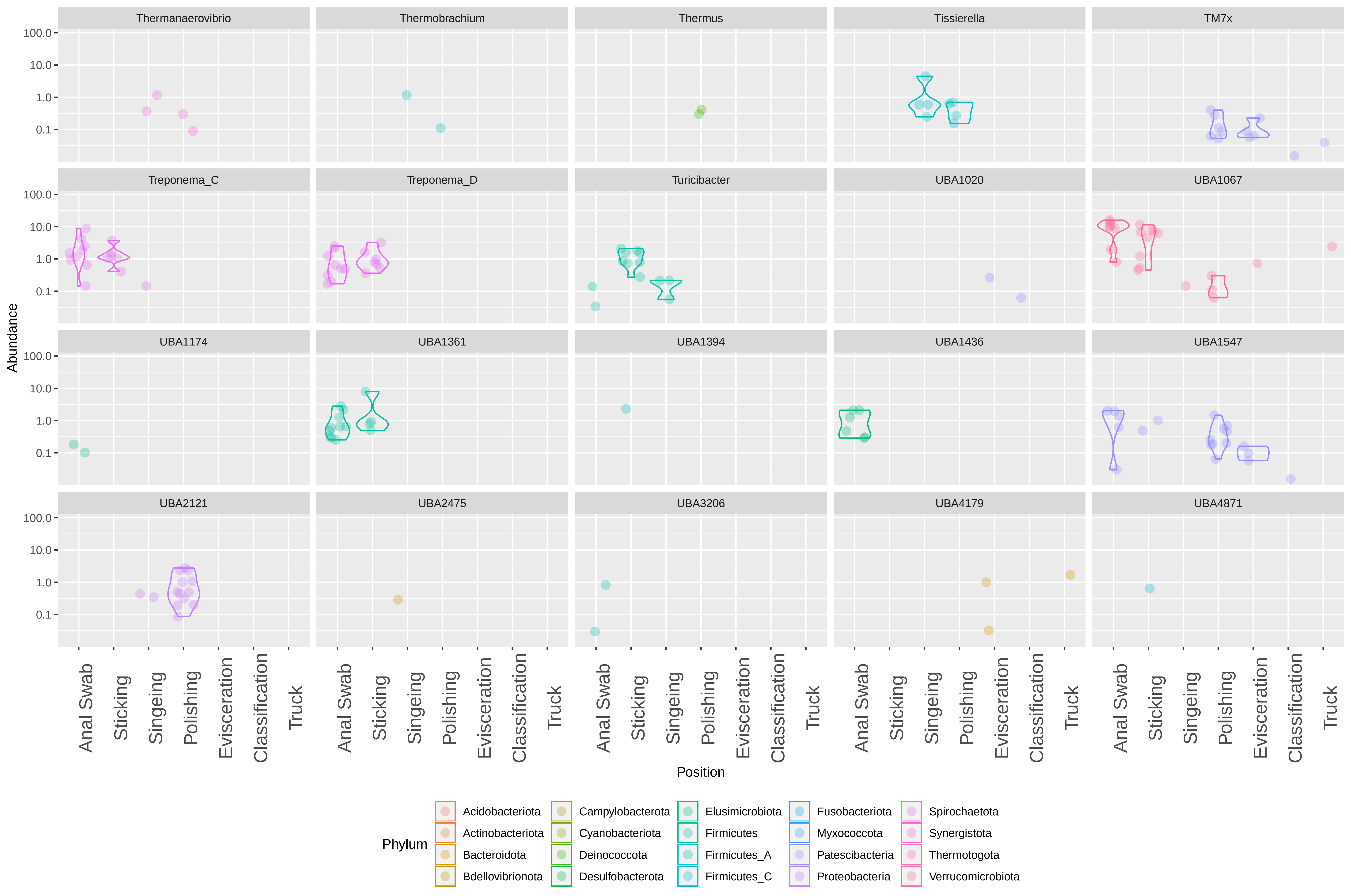

Thermanaerovibrio
Thermobrachium
Thermus
Tissierella
TM7x
Treponema_C
Treponema_D
Turicibacter
UBA1020
UBA1067
UBA1174
UBA1361
UBA1394
UBA1436
UBA1547
UBA2121
UBA2475
UBA3206
UBA4179
UBA4871
100.0
10.0
1.0
0.1
Abundance
Anal Swab
Sticking
Singeing
Polishing
Evisceration
Classification
Truck
Position
Phylum
Acidobacteriota
Actinobacteriota
Bacteroidota
Bdellovibrionota
Campylobacterota
Cyanobacteriota
Deinococcota
Desulfobacterota
Elusimicrobiota
Firmicutes
Firmicutes_A
Firmicutes_C
Fusobacteriota
Myxococcota
Patescibacteria
Proteobacteria
Spirochaetota
Synergistota
Thermotogota
Verrucomicrobiota

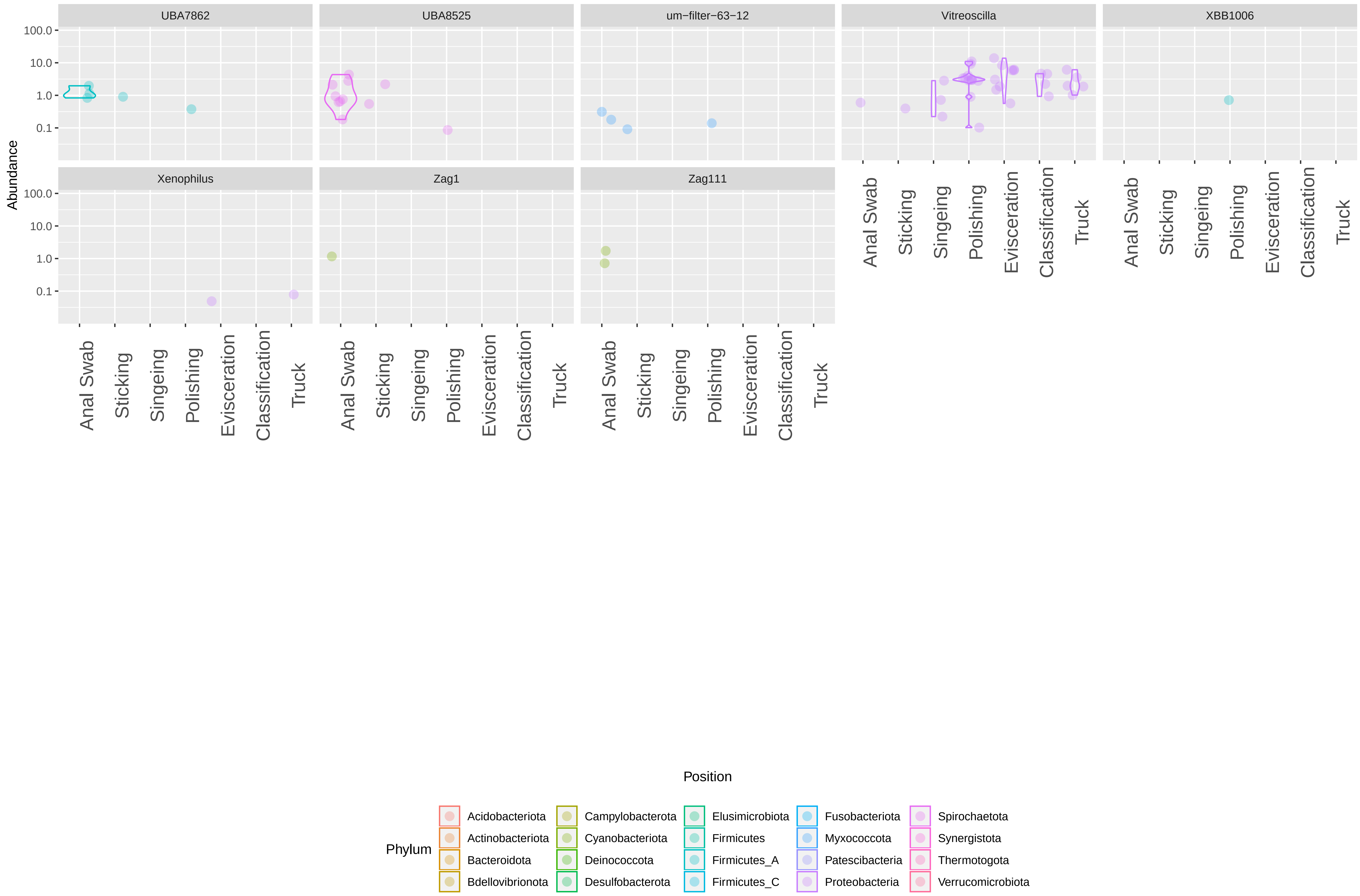
UBA7862
UBA8525
um−filter−63−12
Vitreoscilla
XBB1006
Xenophilus
Zag1
Zag111
Abundance
100.0
10.0
1.0
0.1
Anal Swab
Sticking
Singeing
Polishing
Evisceration
Classification
Truck
Position
Phylum
Acidobacteriota
Actinobacteriota
Bacteroidota
Bdellovibrionota
Campylobacterota
Cyanobacteriota
Deinococcota
Desulfobacterota
Elusimicrobiota
Firmicutes
Firmicutes_A
Firmicutes_C
Fusobacteriota
Myxococcota
Patescibacteria
Proteobacteria
Spirochaetota
Synergistota
Thermotogota
Verrucomicrobiota
