## Supplementary material for "The sources and transmission routes of microbial populations throughout a meat processing facility": Relative abundances of all detected genera across all surface positions

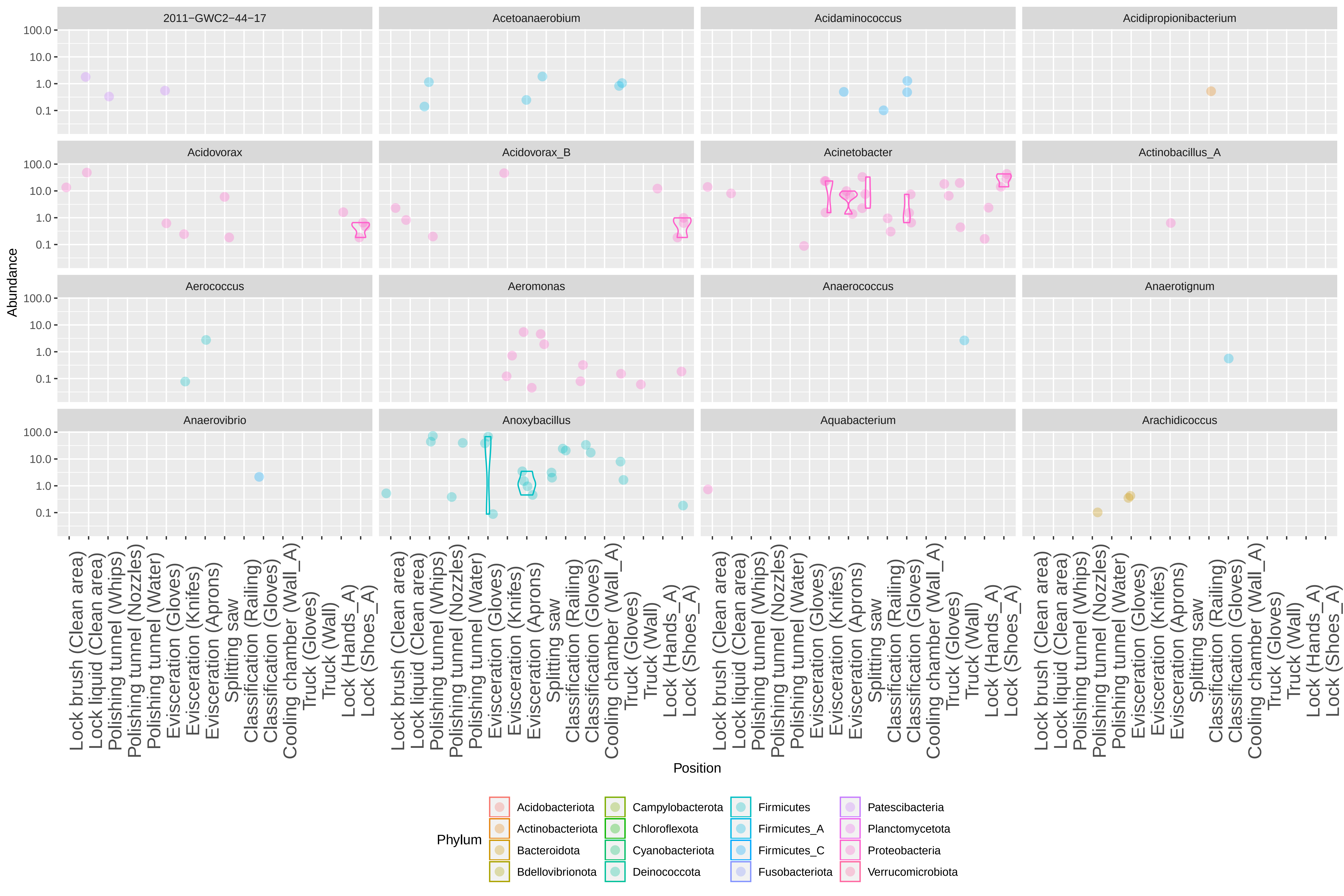
2011−GWC2−44−17
Acetoanaerobium
Acidaminococcus
Acidipropionibacterium
Acidovorax
Acidovorax_B
Acinetobacter
Actinobacillus_A
Aerococcus
Aeromonas
Anaerococcus
Anaerotignum
Anaerovibrio
Anoxybacillus
Aquabacterium
Arachidicoccus
Abundance
100.0
10.0
1.0
0.1
Lock brush (Clean area)
Lock liquid (Clean area)
Polishing tunnel (Whips)
Polishing tunnel (Nozzles)
Polishing tunnel (Water)
Evisceration (Gloves)
Evisceration (Knifes)
Evisceration (Aprons)
Splitting saw
Classification (Railing)
Classification (Gloves)
Cooling chamber (Wall_A)
Truck (Gloves)
Truck (Wall)
Lock (Hands_A)
Lock (Shoes_A)
Position
Phylum
Acidobacteriota
Actinobacteriota
Bacteroidota
Bdellovibrionota
Campylobacterota
Chloroflexota
Cyanobacteriota
Deinococcota
Firmicutes
Firmicutes_A
Firmicutes_C
Fusobacteriota
Patescibacteria
Planctomycetota
Proteobacteria
Verrucomicrobiota

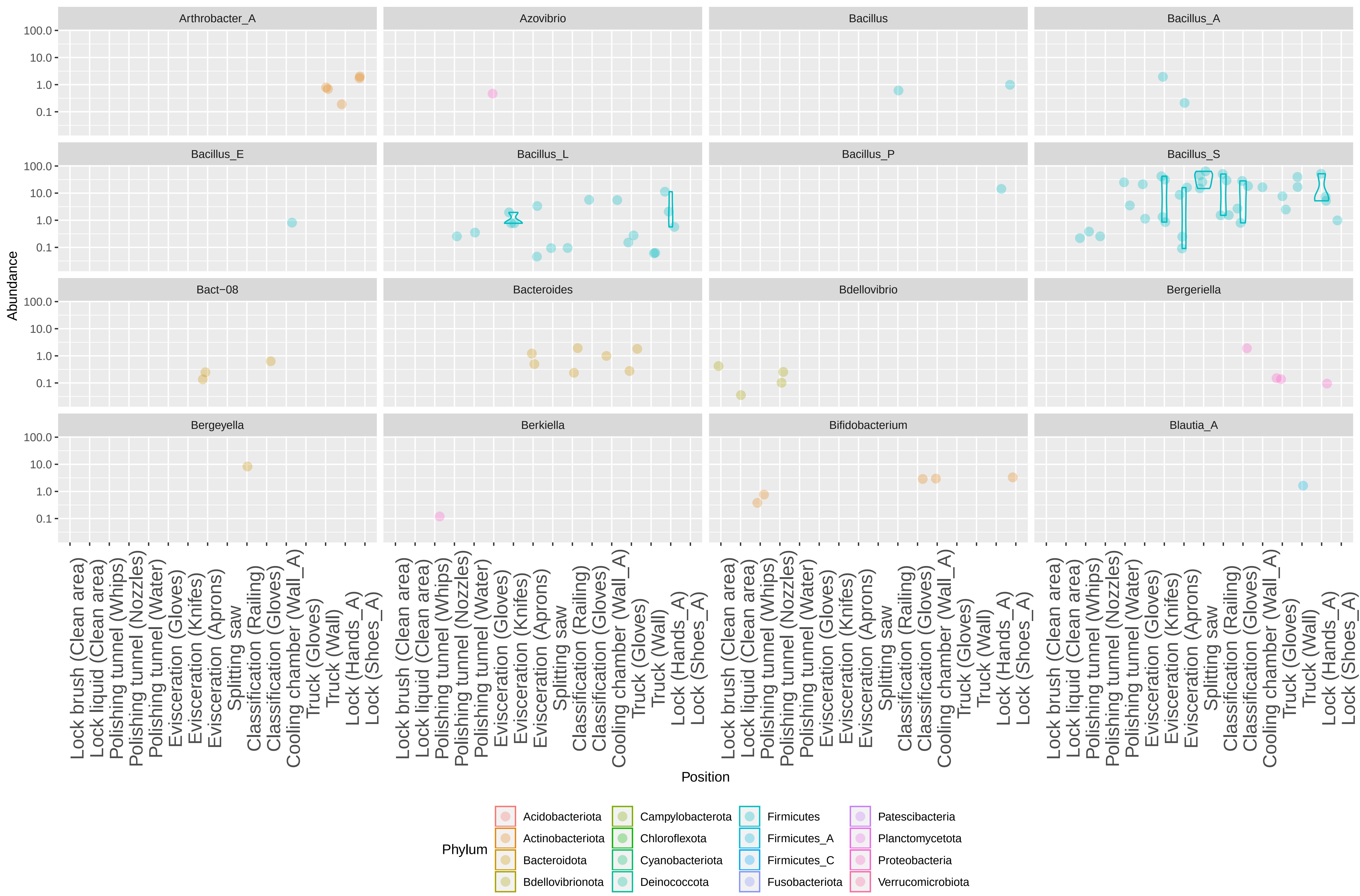

Arthrobacter_A
Azovibrio
Bacillus
Bacillus_A
Bacillus_E
Bacillus_L
Bacillus_P
Bacillus_S
Bact−08
Bacteroides
Bdellovibrio
Bergeriella
Bergeyella
Berkiella
Bifidobacterium
Blautia_A
Abundance
100.0
10.0
1.0
0.1
Lock brush (Clean area)
Lock liquid (Clean area)
Polishing tunnel (Whips)
Polishing tunnel (Nozzles)
Polishing tunnel (Water)
Evisceration (Gloves)
Evisceration (Knifes)
Evisceration (Aprons)
Splitting saw
Classification (Railing)
Classification (Gloves)
Cooling chamber (Wall_A)
Truck (Gloves)
Truck (Wall)
Lock (Hands_A)
Lock (Shoes_A)
Position
Phylum
Acidobacteriota
Actinobacteriota
Bacteroidota
Bdellovibrionota
Campylobacterota
Chloroflexota
Cyanobacteriota
Deinococcota
Firmicutes
Firmicutes_A
Firmicutes_C
Fusobacteriota
Patescibacteria
Planctomycetota
Proteobacteria
Verrucomicrobiota

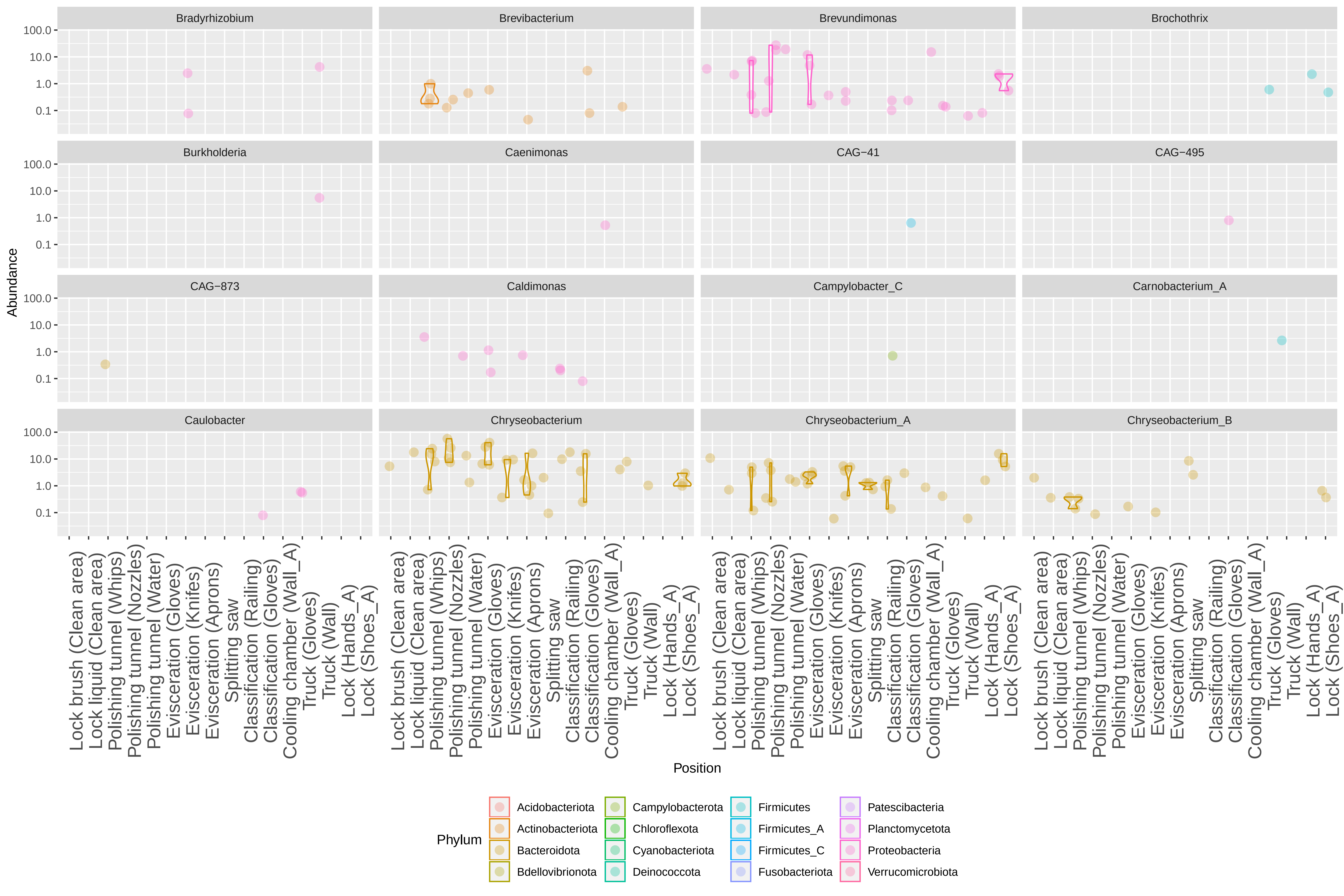

Bradyrhizobium
Brevibacterium
Brevundimonas
Brochothrix
Burkholderia
Caenimonas
CAG−41
CAG−495
CAG−873
Caldimonas
Campylobacter_C
Carnobacterium_A
Caulobacter
Chryseobacterium
Chryseobacterium_A
Chryseobacterium_B
Abundance
100.0
10.0
1.0
0.1
Lock brush (Clean area)
Lock liquid (Clean area)
Polishing tunnel (Whips)
Polishing tunnel (Nozzles)
Polishing tunnel (Water)
Evisceration (Gloves)
Evisceration (Knifes)
Evisceration (Aprons)
Splitting saw
Classification (Railing)
Classification (Gloves)
Cooling chamber (Wall_A)
Truck (Gloves)
Truck (Wall)
Lock (Hands_A)
Lock (Shoes_A)
Position
Phylum
Acidobacteriota
Actinobacteriota
Bacteroidota
Bdellovibrionota
Campylobacterota
Chloroflexota
Cyanobacteriota
Deinococcota
Firmicutes
Firmicutes_A
Firmicutes_C
Fusobacteriota
Patescibacteria
Planctomycetota
Proteobacteria
Verrucomicrobiota

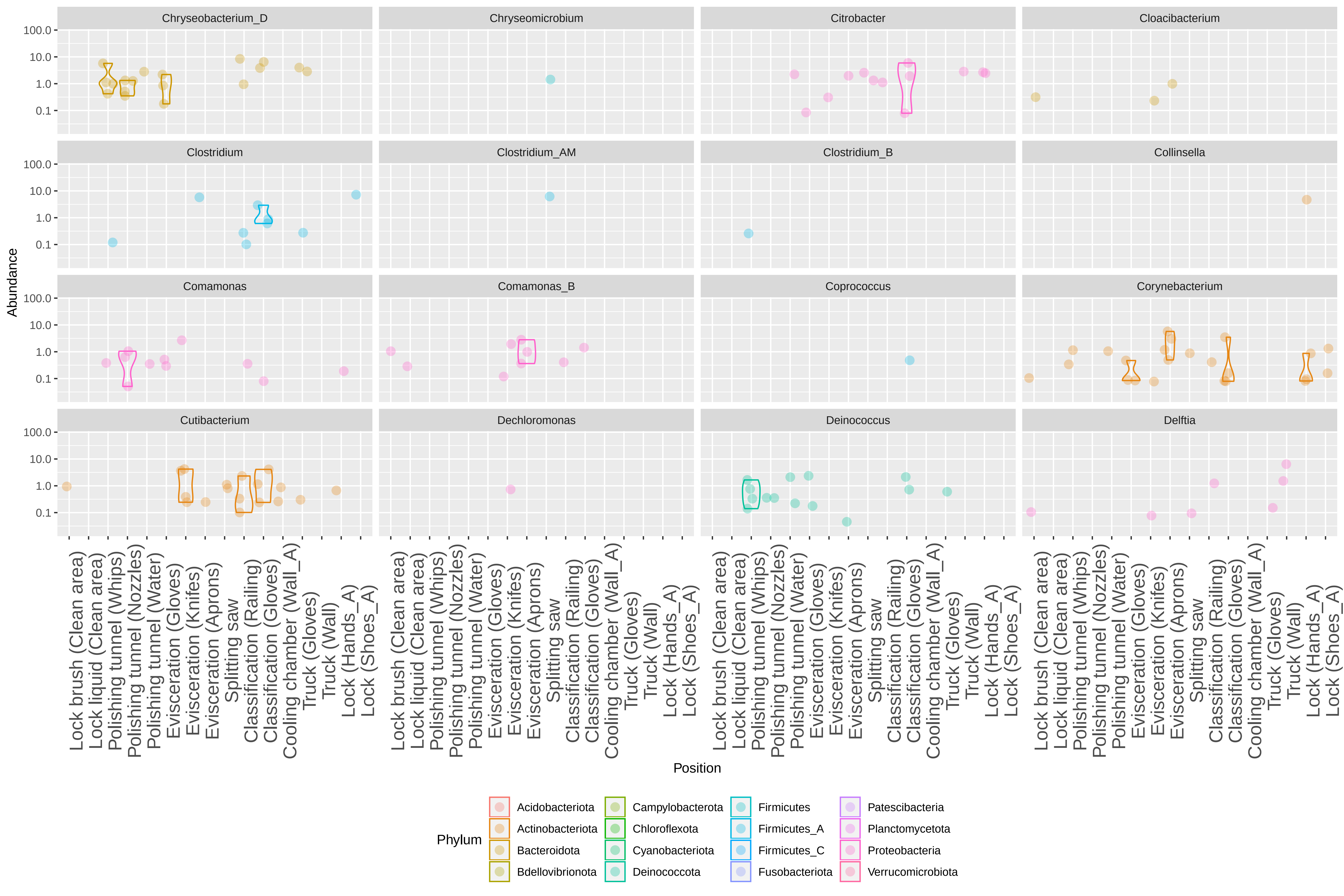

Chryseobacterium_D
Chryseomicrobium
Citrobacter
Cloacibacterium
Clostridium
Clostridium_AM
Clostridium_B
Collinsella
Comamonas
Comamonas_B
Coprococcus
Corynebacterium
Cutibacterium
Dechloromonas
Deinococcus
Delftia
Abundance
100.0
10.0
1.0
0.1
Position
Lock brush (Clean area)
Lock liquid (Clean area)
Polishing tunnel (Whips)
Polishing tunnel (Nozzles)
Polishing tunnel (Water)
Evisceration (Gloves)
Evisceration (Knifes)
Evisceration (Aprons)
Splitting saw
Classification (Railing)
Classification (Gloves)
Cooling chamber (Wall_A)
Truck (Gloves)
Truck (Wall)
Lock (Hands_A)
Lock (Shoes_A)
Phylum
Acidobacteriota
Actinobacteriota
Bacteroidota
Bdellovibrionota
Campylobacterota
Chloroflexota
Cyanobacteriota
Deinococcota
Firmicutes
Firmicutes_A
Firmicutes_C
Fusobacteriota
Patescibacteria
Planctomycetota
Proteobacteria
Verrucomicrobiota

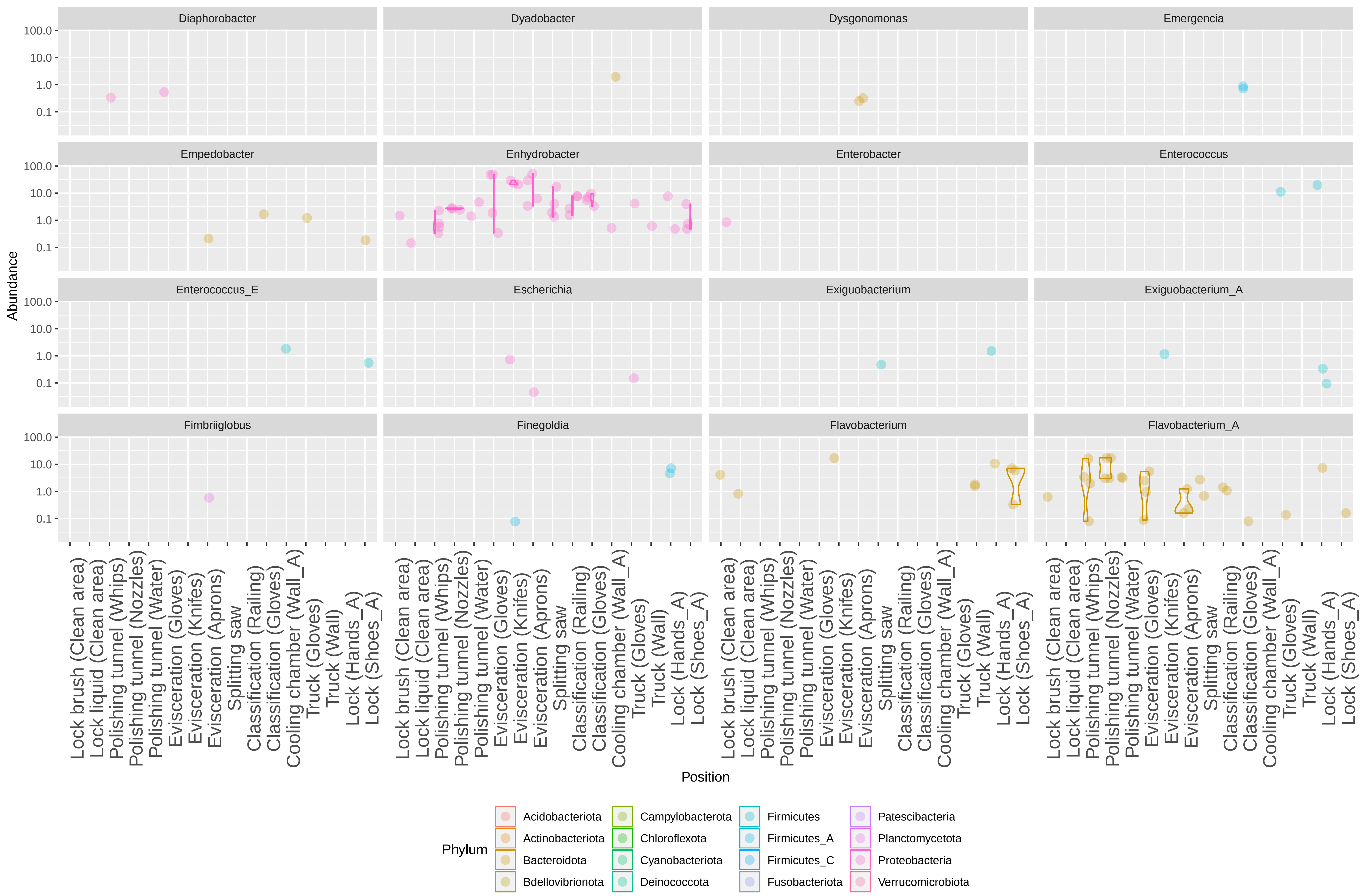

Diaphorobacter
Dyadobacter
Dysgonomonas
Emergencia
Empedobacter
Enhydrobacter
Enterobacter
Enterococcus
Enterococcus_E
Escherichia
Exiguobacterium
Exiguobacterium_A
Fimbriiglobus
Finegoldia
Flavobacterium
Flavobacterium_A
Abundance
100.0
10.0
1.0
0.1
Lock brush (Clean area)
Lock liquid (Clean area)
Polishing tunnel (Whips)
Polishing tunnel (Nozzles)
Polishing tunnel (Water)
Evisceration (Gloves)
Evisceration (Knifes)
Evisceration (Aprons)
Splitting saw
Classification (Railing)
Classification (Gloves)
Cooling chamber (Wall_A)
Truck (Gloves)
Truck (Wall)
Lock (Hands_A)
Lock (Shoes_A)
Position
Phylum
Acidobacteriota
Actinobacteriota
Bacteroidota
Bdellovibrionota
Campylobacterota
Chloroflexota
Cyanobacteriota
Deinococcota
Firmicutes
Firmicutes_A
Firmicutes_C
Fusobacteriota
Patescibacteria
Planctomycetota
Proteobacteria
Verrucomicrobiota

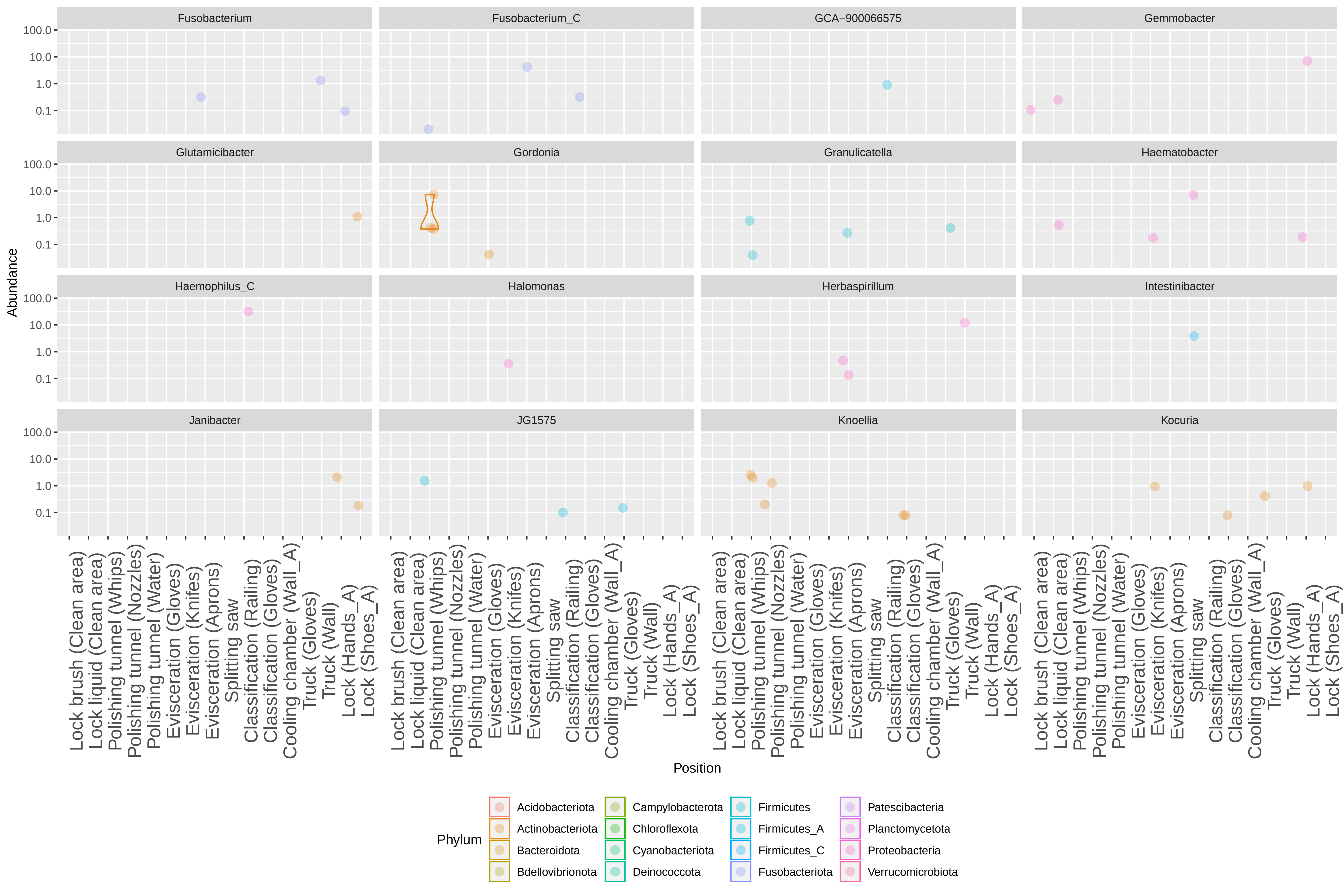

Fusobacterium
Fusobacterium_C
GCA−900066575
Gemmobacter
Glutamicibacter
Gordonia
Granulicatella
Haematobacter
Haemophilus_C
Halomonas
Herbaspirillum
Intestinibacter
Janibacter
JG1575
Knoellia
Kocuria
Abundance
100.0
10.0
1.0
0.1
Position
Lock brush (Clean area)
Lock liquid (Clean area)
Polishing tunnel (Whips)
Polishing tunnel (Nozzles)
Polishing tunnel (Water)
Evisceration (Gloves)
Evisceration (Knifes)
Evisceration (Aprons)
Splitting saw
Classification (Railing)
Classification (Gloves)
Cooling chamber (Wall_A)
Truck (Gloves)
Truck (Wall)
Lock (Hands_A)
Lock (Shoes_A)
Phylum
Acidobacteriota
Actinobacteriota
Bacteroidota
Bdellovibrionota
Campylobacterota
Chloroflexota
Cyanobacteriota
Deinococcota
Firmicutes
Firmicutes_A
Firmicutes_C
Fusobacteriota
Patescibacteria
Planctomycetota
Proteobacteria
Verrucomicrobiota

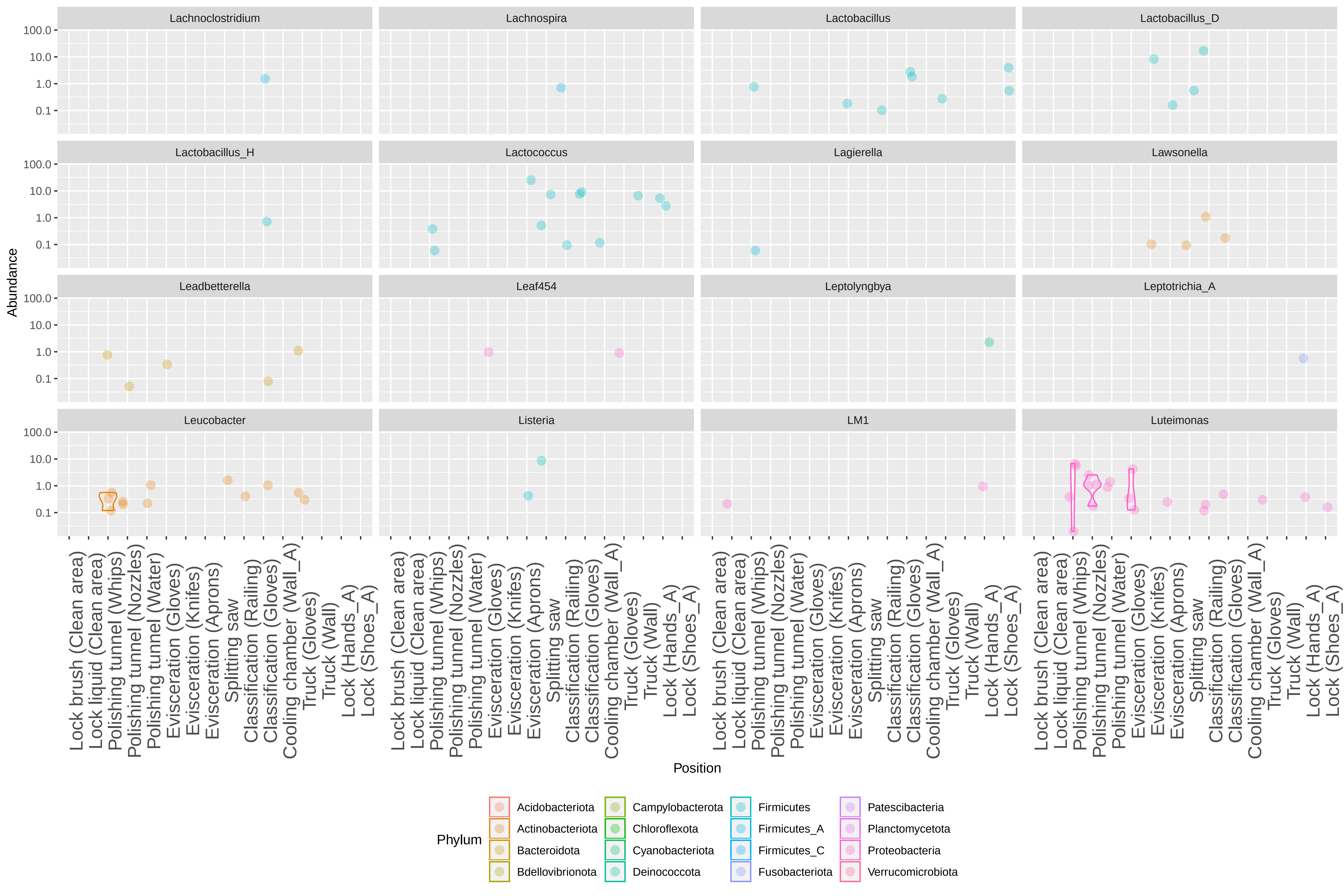

Lachnoclostridium
Lachnospira
Lactobacillus
Lactobacillus_D
Lactobacillus_H
Lactococcus
Lagierella
Lawsonella
Leadbetterella
Leaf454
Leptolyngbya
Leptotrichia_A
Leucobacter
Listeria
LM1
Luteimonas
Abundance
100.0
10.0
1.0
0.1
Lock brush (Clean area)
Lock liquid (Clean area)
Polishing tunnel (Whips)
Polishing tunnel (Nozzles)
Polishing tunnel (Water)
Evisceration (Gloves)
Evisceration (Knifes)
Evisceration (Aprons)
Splitting saw
Classification (Railing)
Classification (Gloves)
Cooling chamber (Wall_A)
Truck (Gloves)
Truck (Wall)
Lock (Hands_A)
Lock (Shoes_A)
Position
Phylum
Acidobacteriota
Actinobacteriota
Bacteroidota
Bdellovibrionota
Campylobacterota
Chloroflexota
Cyanobacteriota
Deinococcota
Firmicutes
Firmicutes_A
Firmicutes_C
Fusobacteriota
Patescibacteria
Planctomycetota
Proteobacteria
Verrucomicrobiota

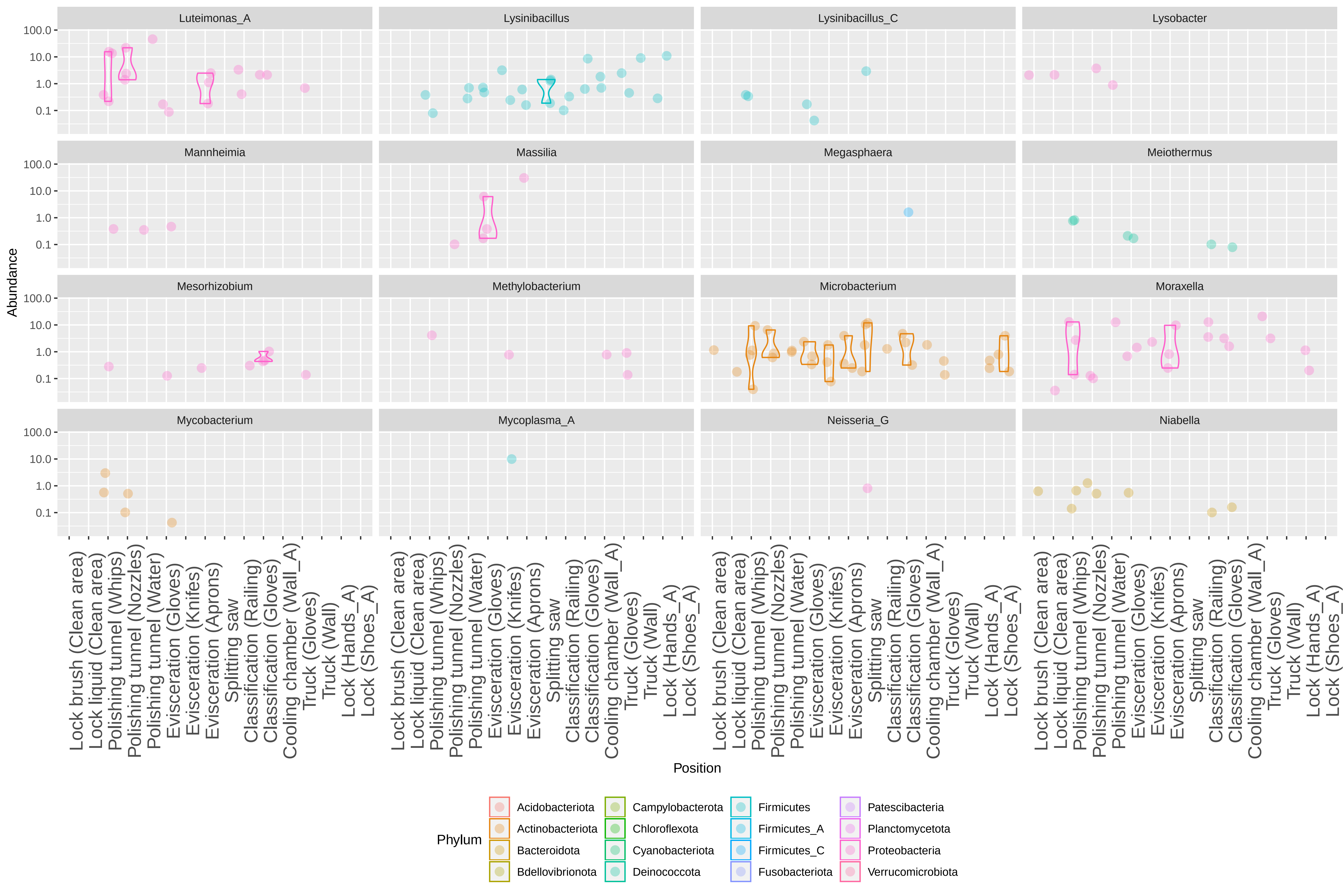

Luteimonas_A
Lysinibacillus
Lysinibacillus_C
Lysobacter
Mannheimia
Massilia
Megasphaera
Meiothermus
Mesorhizobium
Methylobacterium
Microbacterium
Moraxella
Mycobacterium
Mycoplasma_A
Neisseria_G
Niabella
Abundance
100.0
10.0
1.0
0.1
Position
Lock brush (Clean area)
Lock liquid (Clean area)
Polishing tunnel (Whips)
Polishing tunnel (Nozzles)
Polishing tunnel (Water)
Evisceration (Gloves)
Evisceration (Knifes)
Evisceration (Aprons)
Splitting saw
Classification (Railing)
Classification (Gloves)
Cooling chamber (Wall_A)
Truck (Gloves)
Truck (Wall)
Lock (Hands_A)
Lock (Shoes_A)
Phylum
Acidobacteriota
Actinobacteriota
Bacteroidota
Bdellovibrionota
Campylobacterota
Chloroflexota
Cyanobacteriota
Deinococcota
Firmicutes
Firmicutes_A
Firmicutes_C
Fusobacteriota
Patescibacteria
Planctomycetota
Proteobacteria
Verrucomicrobiota

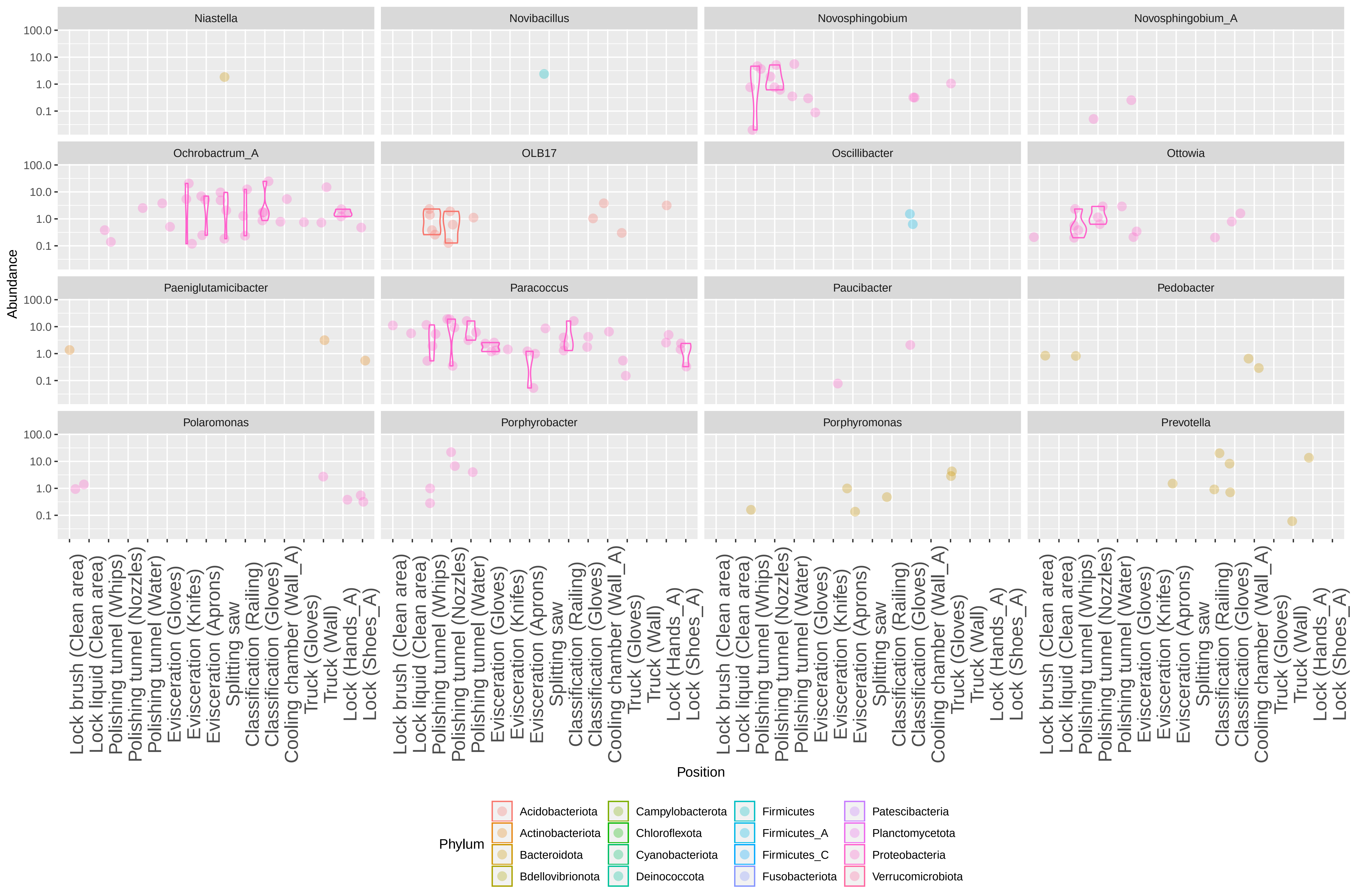

Niastella
Novibacillus
Novosphingobium
Novosphingobium_A
Ochrobactrum_A
OLB17
Oscillibacter
Ottowia
Paeniglutamicibacter
Paracoccus
Paucibacter
Pedobacter
Polaromonas
Porphyrobacter
Porphyromonas
Prevotella
Abundance
100.0
10.0
1.0
0.1
Lock brush (Clean area)
Lock liquid (Clean area)
Polishing tunnel (Whips)
Polishing tunnel (Nozzles)
Polishing tunnel (Water)
Evisceration (Gloves)
Evisceration (Knifes)
Evisceration (Aprons)
Splitting saw
Classification (Railing)
Classification (Gloves)
Cooling chamber (Wall_A)
Truck (Gloves)
Truck (Wall)
Lock (Hands_A)
Lock (Shoes_A)
Position
Phylum
Acidobacteriota
Actinobacteriota
Bacteroidota
Bdellovibrionota
Campylobacterota
Chloroflexota
Cyanobacteriota
Deinococcota
Firmicutes
Firmicutes_A
Firmicutes_C
Fusobacteriota
Patescibacteria
Planctomycetota
Proteobacteria
Verrucomicrobiota

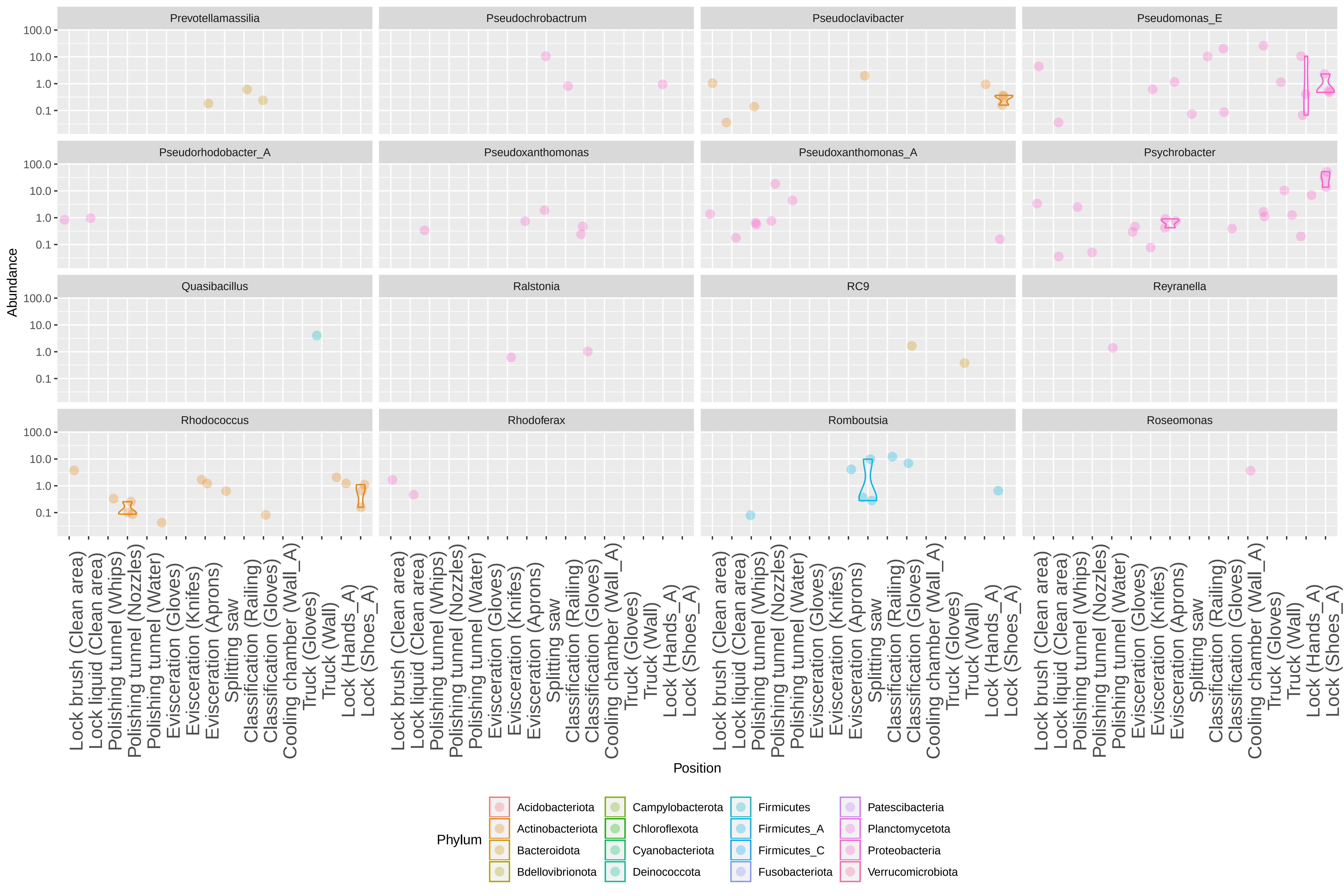

Prevotellamassilia
Pseudochrobactrum
Pseudoclavibacter
Pseudomonas_E
Pseudorhodobacter_A
Pseudoxanthomonas
Pseudoxanthomonas_A
Psychrobacter
Quasibacillus
Ralstonia
RC9
Reyranella
Rhodococcus
Rhodoferax
Romboutsia
Roseomonas
Abundance
100.0
10.0
1.0
0.1
Lock brush (Clean area)
Lock liquid (Clean area)
Polishing tunnel (Whips)
Polishing tunnel (Nozzles)
Polishing tunnel (Water)
Evisceration (Gloves)
Evisceration (Knifes)
Evisceration (Aprons)
Splitting saw
Classification (Railing)
Classification (Gloves)
Cooling chamber (Wall_A)
Truck (Gloves)
Truck (Wall)
Lock (Hands_A)
Lock (Shoes_A)
Position
Phylum
Acidobacteriota
Actinobacteriota
Bacteroidota
Bdellovibrionota
Campylobacterota
Chloroflexota
Cyanobacteriota
Deinococcota
Firmicutes
Firmicutes_A
Firmicutes_C
Fusobacteriota
Patescibacteria
Planctomycetota
Proteobacteria
Verrucomicrobiota

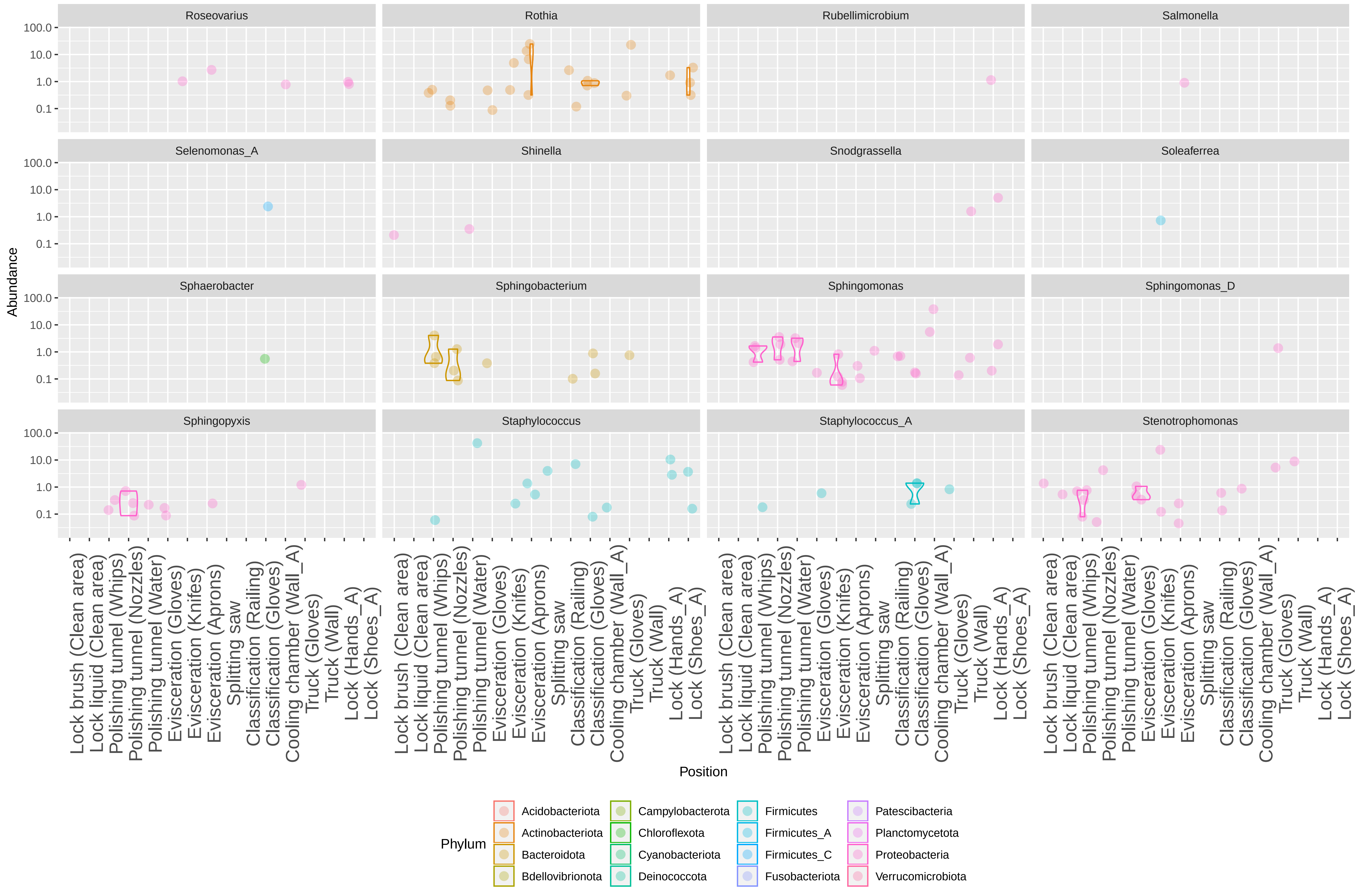

Roseovarius
Rothia
Rubellimicrobium
Salmonella
Selenomonas_A
Shinella
Snodgrassella
Soleaferrea
Sphaerobacter
Sphingobacterium
Sphingomonas
Sphingomonas_D
Sphingopyxis
Staphylococcus
Staphylococcus_A
Stenotrophomonas
100.0
10.0
1.0
0.1
Abundance
Lock brush (Clean area)
Lock liquid (Clean area)
Polishing tunnel (Whips)
Polishing tunnel (Nozzles)
Polishing tunnel (Water)
Evisceration (Gloves)
Evisceration (Knifes)
Evisceration (Aprons)
Splitting saw
Classification (Railing)
Classification (Gloves)
Cooling chamber (Wall_A)
Truck (Gloves)
Truck (Wall)
Lock (Hands_A)
Lock (Shoes_A)
Position
Phylum
Acidobacteriota
Actinobacteriota
Bacteroidota
Bdellovibrionota
Campylobacterota
Chloroflexota
Cyanobacteriota
Deinococcota
Firmicutes
Firmicutes_A
Firmicutes_C
Fusobacteriota
Patescibacteria
Planctomycetota
Proteobacteria
Verrucomicrobiota

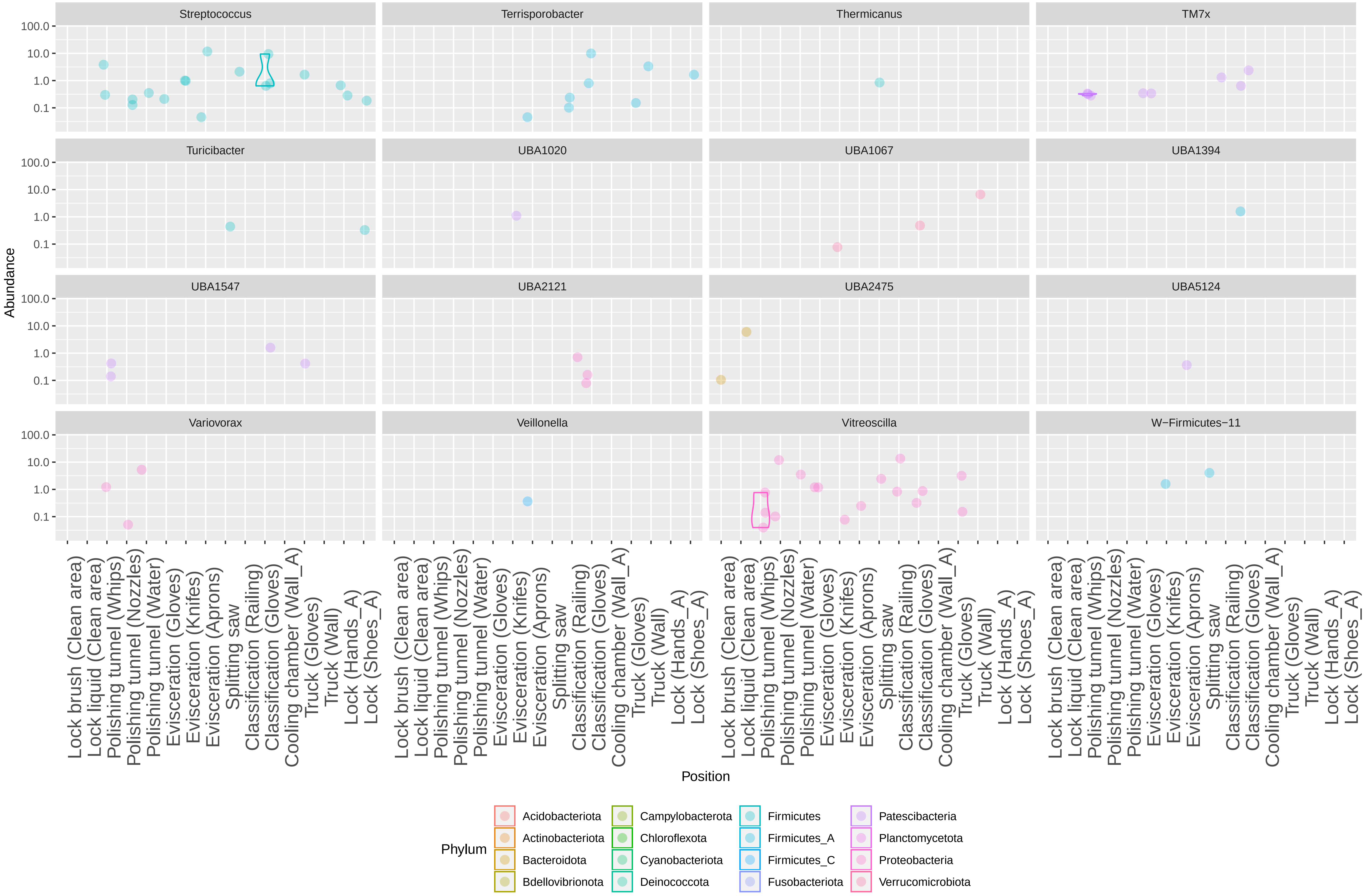
Streptococcus
Terrisporobacter
Thermicanus
TM7x
Turicibacter
UBA1020
UBA1067
UBA1394
UBA1547
UBA2121
UBA2475
UBA5124
Variovorax
Veillonella
Vitreoscilla
W−Firmicutes−11
Abundance
100.0
10.0
1.0
0.1
Position
Lock brush (Clean area)
Lock liquid (Clean area)
Polishing tunnel (Whips)
Polishing tunnel (Nozzles)
Polishing tunnel (Water)
Evisceration (Gloves)
Evisceration (Knifes)
Evisceration (Aprons)
Splitting saw
Classification (Railing)
Classification (Gloves)
Cooling chamber (Wall_A)
Truck (Gloves)
Truck (Wall)
Lock (Hands_A)
Lock (Shoes_A)
Phylum
Acidobacteriota
Actinobacteriota
Bacteroidota
Bdellovibrionota
Campylobacterota
Chloroflexota
Cyanobacteriota
Deinococcota
Firmicutes
Firmicutes_A
Firmicutes_C
Fusobacteriota
Patescibacteria
Planctomycetota
Proteobacteria
Verrucomicrobiota

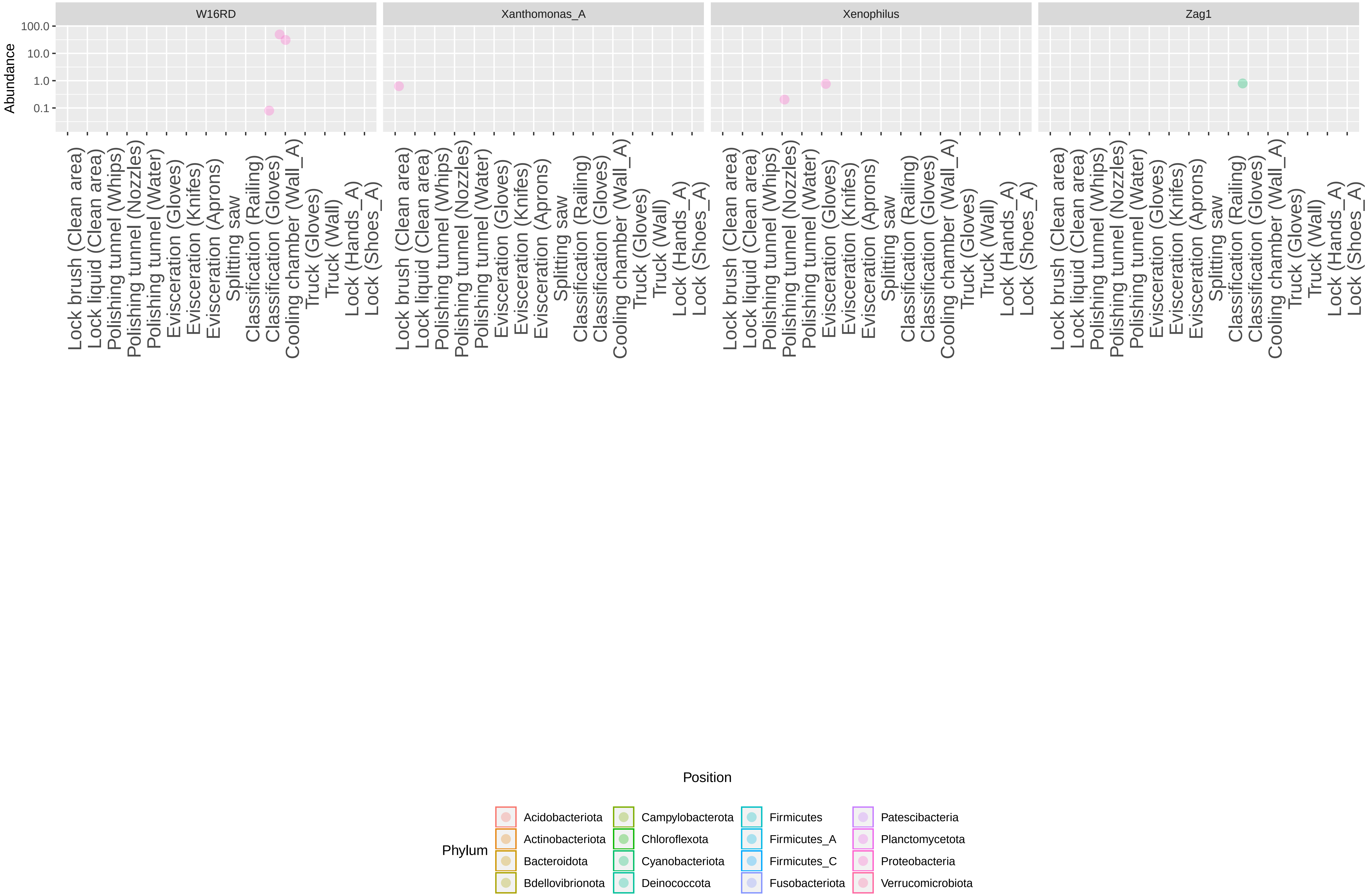

W16RD
Xanthomonas_A
Xenophilus
Zag1
Abundance
100.0
10.0
1.0
0.1
Lock brush (Clean area)
Lock liquid (Clean area)
Polishing tunnel (Whips)
Polishing tunnel (Nozzles)
Polishing tunnel (Water)
Evisceration (Gloves)
Evisceration (Knifes)
Evisceration (Aprons)
Splitting saw
Classification (Railing)
Classification (Gloves)
Cooling chamber (Wall_A)
Truck (Gloves)
Truck (Wall)
Lock (Hands_A)
Lock (Shoes_A)
Position
Phylum
Acidobacteriota
Actinobacteriota
Bacteroidota
Bdellovibrionota
Campylobacterota
Chloroflexota
Cyanobacteriota
Deinococcota
Firmicutes
Firmicutes_A
Firmicutes_C
Fusobacteriota
Patescibacteria
Planctomycetota
Proteobacteria
Verrucomicrobiota
